# Genetic evidence for a regulated cysteine protease catalytic triad in LegA7, a *Legionella pneumophila* protein that impinges on a stress response pathway

**DOI:** 10.1101/2024.03.17.585421

**Authors:** Dar Hershkovitz, Emy J. Chen, Alexander W. Ensminger, Aisling S. Dugan, Kaleigh T. Conway, Alex C. Joyce, Gil Segal, Ralph R. Isberg

## Abstract

*Legionella pneumophila* grows within membrane-bound vacuoles in phylogenetically diverse hosts. Intracellular growth requires the function of the Icm/Dot type-IVb secretion system, which translocates more than 300 proteins into host cells. A screen was performed to identify *L. pneumophila* proteins that stimulate MAPK activation, using Icm/Dot translocated proteins ectopically expressed in mammalian cells. In parallel, a second screen was performed to identify *L. pneumophila* proteins expressed in yeast that cause growth inhibition in MAPK pathway-stimulatory high osmolarity medium. LegA7 was shared in both screens, a protein predicted to be a member of the bacterial cysteine protease family that has five carboxyl-terminal ankyrin repeats. Three conserved residues in the predicted catalytic triad of LegA7 were mutated. These mutations abolished the ability of LegA7 to inhibit yeast growth. To identify other residues important for LegA7 function, a generalizable selection strategy in yeast was devised to isolate mutants that have lost function and no longer cause growth inhibition on high osmolarity medium. Mutations were isolated in the two carboxyl-terminal ankyrin repeats, as well as an inter-domain region located between the cysteine protease domain and the ankyrin repeats. These mutations were predicted by AlphaFold modeling to localize to the face opposite from the catalytic site, arguing that they interfere with the positive regulation of the catalytic activity. Based on our data, we present a model in which LegA7 harbors a cysteine protease domain with an inter-domain and two carboxyl-terminal ankyrin repeat regions that modulate the function of the catalytic domain.

## INTRODUCTION

*Legionella pneumophila* is a Gram-negative, facultative, intracellular bacterium and the causative agent of Legionnaires’ disease (1, 2), which presents with either atypical pneumonia or flu-like symptoms, such as cough, fever, and chills. Individuals who are immunocompromised, smokers, or have chronic lung disease are at increased risk for disease (3). Disease is caused by inhalation or aspiration of aerosolized bacteria from freshwater sources, which are then internalized by alveolar macrophages (4). It is thought that amoebae growing in freshwater sources, such as *Acanthamoeba castellanii* and *Hartmannella vermiformis*, are the reservoirs for the bacteria that cause disease in humans (5, 6). Selection for disease, thus, is thought to occur entirely in nonmammalian hosts.

Inside host cells, *L. pneumophila* resides and replicates in a membrane-bound vacuole that avoids fusion with late endosomes and lysosomes (7, 8). Both tubular endoplasmic reticulum (ER) and secretory vesicle-derived material is recruited to the surface of the *Legionella-*containing vacuole (LCV), resulting in a compartment surrounded by ER (7, 9–13). Central to *L. pneumophila* pathogenesis is the Icm/Dot type IVb secretion system which is critical for the formation of the LCV (14, 15). Each *L. pneumophila* isolate injects over 300 different bacterial proteins through this secretion system into host cells (16–19). The total number of these translocated effectors identified in the *Legionella* pangenome is staggering, as machine learning strategies have identified at least 18,000 such proteins among isolates from more than 80 *Legionella* species (20, 21).

The roles of individual Icm/Dot translocated substrates (IDTS) in promoting replication and vacuole formation have been difficult to elucidate because individual deletions of most substrates of the *L. pneumophila* genome have no consequence on intracellular growth of the bacterium. This lack of defect is thought to be due to functional redundancy, such that multiple translocated substrates target parallel pathways in the host cell or are able to complete a pathway independently of each other (22). Recent work indicates that IDTS expansion and redundancy can be partly explained by selection for growth in multiple poorly related amoebal hosts (23), and temporal overlap in the execution of individual protein activities during intracellular growth (24, 25).

Previous work has shown that mitogen-activated protein kinases (MAPKs) are modulated during *L. pneumophila* infection. Upon *L. pneumophila* challenge in amoebae, a MAPK response interferes with *L. pneumophila* intracellular growth (26), consistent with an evolutionarily conserved pathway that controls *L. pneumophila* growth. In addition, *L. pneumophila* challenge of mammalian cells leads to a cytokine response via NFκB and MAPK signaling (27, 28). Finally, several IDTS have been shown to interfere with host cell protein synthesis and activate the MAPK response (28, 29), indicating that MAPKs may be common targets of *L. pneumophila* virulence-associated proteins.

The MAPK family of serine/threonine kinases is involved in directing cellular responses to a diverse array of stimuli such as growth factors, mitogens, osmotic and oxidative stress, and inflammatory cytokines (30–32). Members regulate proliferation, cell division, differentiation, apoptosis, inflammation, growth, and gene expression (30–32). The mammalian MAP kinase family includes the extracellular signal-related kinases (ERK1 and ERK2), c-Jun NH_2_-terminal kinases (JNK1, JNK2, and JNK3), and p38 proteins (p38α, p38β, p38γ and p38δ). MAPKs are activated by phosphorylation by specific MAPK kinases, MAP2Ks. The MAP2Ks are, in turn, activated by MAP2K kinases (MAP3Ks), which receive upstream signals such as growth factors binding their respective cell-surface receptors, or chemical or physical stresses in the extracellular environment (31). MAPKs are targets of numerous effectors translocated into host cells by diverse bacterial pathogens (33).

In the following study, complementary strategies were taken to identify *L. pneumophila* proteins that promote stresses associated with the MAPK response. A putative member of the bacterial cysteine protease family having carboxyl terminal ankyrin repeats was identified. We provide evidence that the function of this effector requires both a conserved catalytic triad and the amino-terminal ankyrin repeats.

## RESULTS

### Identification of Icm/Dot substrates that cause elevated phosphorylation of SAPK/JNK in mammalian cells

Proteins had been previously identified that were able to activate an NFκB-regulated promoter by expressing a bank of *L. pneumophila* Icm/Dot translocated substrate (IDTS) genes in mammalian cells (34). We took a similar approach to identify bacterial proteins that activate MAPKs, because a variety of studies show that MAPKs either modulate *L. pneumophila* intracellular growth or respond to specific IDTS (26–29). To identify IDTS that activate mammalian MAPKs, we constructed a bank of 257 known and putative IDTS genes fused to *gfp* in a mammalian expression vector, expanding our previously constructed library (Materials and Methods; (34)). This bank was then transfected into HEK293T cells to identify *L. pneumophila* proteins that could activate MAPK cascades, as determined by increased phosphorylation of either ERK, p38, or SAPK/JNK relative to the empty vector control, using phosphorylation-specific antibodies (Material and Methods). As each of the MAPKs gave similar levels of activation in response to the IDTS, repetitions used only SAPK/JNK phosphorylation as the readout (Table 1).

**TABLE 1.** Identification of Icm/Dot translocated substrates that result in increased phospho-JNK levels in mammalian cells.

| Lpg number <sup>a</sup> | Gene | Relative SAPK/JNK activation | MAD score |
| --- | --- | --- | --- |
| lpg0234 | *sidE* | 15.00 | 48.60 |
| lpg2157 | *sdeA* | 5.61 | 15.65 |
| lpg2156 | *sdeB* | 5.19 | 14.18 |
| lpg0059 | *ceg2* | 4.87 | 13.05 |
| lpg1701 | *legC3* | 4.00 | 10.00 |
| lpg2831 | *vipD* | 3.34 | 7.68 |
| lpg0695 | *legA8* | 3.30 | 7.54 |
| lpg0080 | *ceg3* | 3.00 | 6.49 |
| lpg2207 |  | 2.76 | 5.65 |
| lpg1960 | *lirA* | 2.63 | 5.19 |
| lpg2147 | *mavC* | 2.50 | 4.74 |
| lpg2153 | *sdeC* | 2.50 | 4.74 |
| lpg1752 | *mavB* | 2.40 | 4.39 |
| lpg0240 | *ceg8* | 2.36 | 4.25 |
| lpg2148 | *mvcA* | 2.35 | 4.21 |
| lpg2498 | *mavJ* | 2.34 | 4.18 |
| lpg2456 | *legA15* | 2.33 | 4.14 |
| lpg0160 | *ravD* | 2.32 | 4.11 |
| lpg1961 |  | 2.24 | 3.82 |
| lpg2862 | *legC8* | 2.24 | 3.82 |
| lpg0403 | *legA7* | 2.23 | 3.79 |
| lpg1488 | *legC5* | 2.22 | 3.75 |
| lpg2420 |  | 2.20 | 3.68 |
| lpg2327 | *ceg6* | 2.20 | 3.68 |

<sup>a</sup> Listed are *L. pneumophila* genes that, when ectopically expressed from the pDEST53 plasmid, result in enhanced JNK phosphorylation relative to empty vector control, using the median absolute deviation (MAD) score $\geq 3.5$ as a cutoff for increased JNK phosphorylation relative to wild type control.

A rank-order table was generated based on the median absolute deviation (MAD; Materials and Methods) of SAPK/JNK phosphorylation for each transfectant relative to empty vector control, and 24 candidates were identified that caused increased levels of SAPK/JNK phosphorylation, based on the Iglewicz and Hoaglin outlier test ((35); MAD >3.5; Table 1). IDTS that resulted in enhanced SAPK/JNK phosphorylation included a number of members of the SidE family of IDTS that catalyze phosphoribosyl-linked ubiquitin modification of targets (13, 36–38), the previously characterized AnkX phosphocholine transferase (39), as well as the Rab5-activated phospholipase VipD (40–42). To focus on a subset of IDTS candidates that affected known MAPK-activated pathways, we performed a screen in yeast to study the MAPK-related response.

### Expression of *legA7* in yeast inhibits osmoadaptation

MAPK activation in response to *L. pneumophila* occurs in highly diverse and evolutionarily unrelated cell types such as mouse macrophages and *Dictyostelium discoideum* amoebae (26–28). To further whittle down the candidates to pursue further, we sought to identify IDTS that either impinge or synergize with the MAPK cascade across species boundaries. We previously constructed a bank of *Saccharomyces cerevisiae* strains harboring ectopically expressed IDTS under the GAL1 promoter control (43), so this bank was used to identify candidate IDTS that modulate the ability of yeast to respond to stresses that activate the MAPK cascade, taking advantage of a simple plate assay. To this end, we characterized IDTS previously demonstrated to cause growth defects in *S. cerevisiae* when expressed in a medium containing galactose to induce expression (43). For most strains expressing IDTS, the defect was dependent on the presence of galactose in the growth medium, consistent with the growth defects being dependent on the transcriptional induction of each of the IDTS genes (Supplemental Dataset 1; Fig. 1).

**Figure 1.**
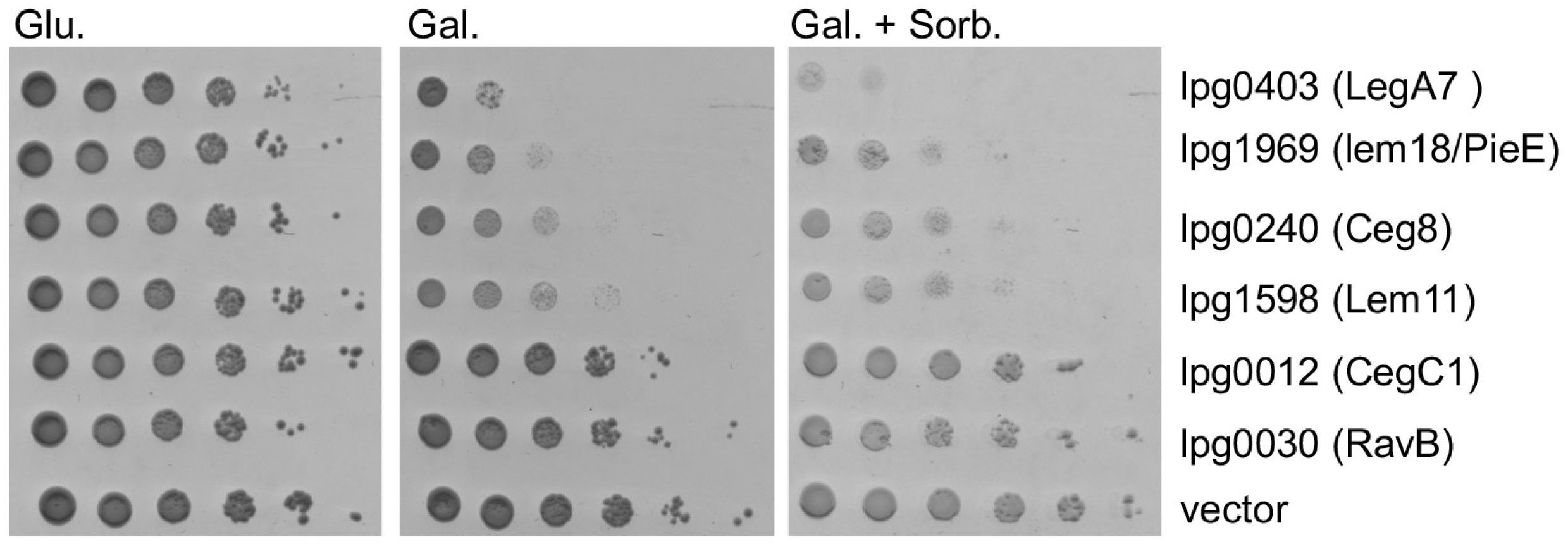
Growth defects caused by *L. pneumophila* Icm/Dot translocated substrates expressed in *S. cerevisiae*. The *L. pneumophila* effectors (indicated on the right) were cloned under the GAL1 promoter and grown on plates containing glucose (Glu), galactose (Gal, inducing conditions), or galactose supplemented with 1 M sorbitol (Gal. + Sorb.) at 30°C, in the wild-type *S. cerevisiae* BY4741. pGREG523 (vector) was used as a negative control. Ten-fold serial dilutions were performed, and the growth inhibition effect was compared to the one of the vector pGREG523 control (vector).

Based on the behavior of the strains plated on galactose-containing medium, many of the *S. cerevisiae* strains harboring IDTS either showed mild growth defects or still had detectable colony formation after serial dilution (Dataset S1; for examples see Fig. 1). We predicted that some of the strains with intermediate phenotypes would have amplified defects on medium that activates a stress pathway requiring MAPK activity for viability. After induction of IDTS expression, such strains should show lower colony formation efficiency in the presence of stress conditions, when compared to the same medium that did not induce this stress.

Wild-type yeast can adapt to a variety of environmental stresses, such as high osmolarity, allowing growth in the presence of high concentrations of a variety of solutes. Growth under these conditions depends on MAPK activation to provide osmoprotection (44). To address whether high osmolarity could potentiate *S. cerevisiae* growth defects caused by the expression of *L. pneumophila* IDTS, a screen was performed to identify osmosensitive strains (Dataset S1). Using the strains that showed depressed CFU formation after galactose induction, we screened for amplification of these defects by plating on a high osmolarity medium containing sorbitol (45–48). Strains were then retained that showed lower CFU efficiency on sorbitol-containing medium in the presence of galactose relative to the identical medium lacking sorbitol. The strain harboring a plasmid encoding *legA7* (lpg0403), which is an uncharacterized IDTS (49), had the lowest colony-forming efficiency on high sorbitol relative to the screened pool (Dataset S1; Fig. 1). *S. cerevisiae* harboring *legA7* showed a reduction in CFU on a medium containing high sorbitol compared to the empty vector control (Fig. 1). No strains harboring IDTS showed enhanced growth in the presence of sorbitol.

### Expression of LegA7 in mammalian cells results in hyperphosphorylation of SAPK/JNK

LegA7, also referred to as AnkZ (50), had been identified as a potential translocated substrate based on the presence of predicted ankyrin repeats in its sequence (49). Consistent with its designation as an IDTS, the protein was shown to contain an Icm/Dot recognition signal that allows the translocation of reporter constructs into mammalian cells (50). Furthermore, transfection with a LegA7-expressing plasmid resulted in high MAPK activation levels (Table 1). To demonstrate that this screen was an accurate representation of the levels of activation by LegA7, HEK293T cells were transfected in triplicate with a plasmid encoding LegA7, as well as two other plasmids encoding IDTS that had caused some level of sorbitol sensitivity to yeast (Dataset S1; lpg0030, and lpg0059). Transfectants harboring the plasmid encoding LegA7 clearly showed enhanced phosphorylation of SAPK/JNK relative to the empty vector control (Fig. 2A; quantitated in 2B), although blotting of the GFP-LegA7 fusion showed that the predominant form of the protein present at steady state was missing the carboxyl-terminal 20 kD based on electrophoretic mobility determination of apparent molecular weights (Fig. 2A, arrow). We believe this result was a consequence of autodegradation (Supplemental Fig. S1; Fig. 5). Therefore, LegA7 induced a MAPK response in mammalian cells in addition to its potentiation of osmotic stress in lower eukaryotes.

**Figure 2.**
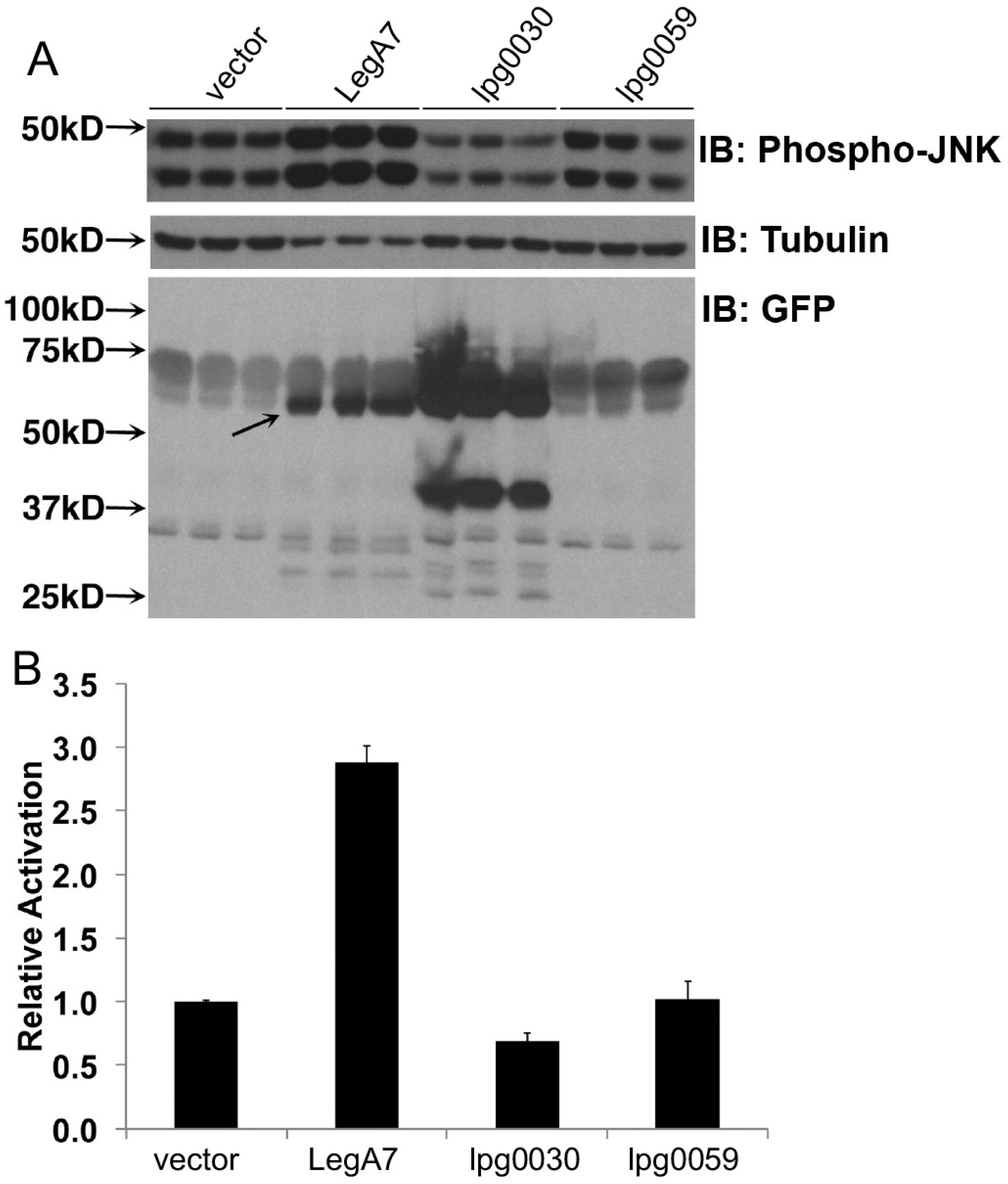
Ectopic expression of LegA7 results in elevated phosphorylation of SAPK/JNK in mammalian cells. **A.** Quantitation of phospho-SAPK/JNK levels in the mammalian cell line HEK293T cells during ectopic expression of LegA7, Lpg0030, and Lpg0059 relative to empty vector control pDEST53 (vector). 40 hours after transfection, extracts were prepared, fractionated, and immunoprobed with anti-phospho-JNK (top panel), α-tubulin (middle panel), and GFP to identify hybrid proteins (bottom panel). Samples were run in triplicate, showing three independent transfections of plasmids into HEK293T. The arrow points to the LegA7 degradation product. Predicted sizes for GFP fusions are: empty vector, 27 kDa; LegA7, 84 kDa; Lpg0030, 62 kDa; Lpg0059 68 kDa. **B.** Increased phosphorylation of LegA7 relative to other IDTS. Data were quantitated by determining phosphorylation levels relative to tubulin loading control (Materials and Methods). Data are the mean of three samples ± S.E.

### The effect of LegA7 on yeast is connected to MAPK pathways

To explore the connection between LegA7 and MAPK-related stresses in yeast, we examined the effect of LegA7 on yeast growth under four stress conditions that are known to activate MAPK pathways. Like Sorbitol, NaCl, and cold temperature (20°C) activate the high-osmolarity glycerol (HOG) pathway that controls the osmoregulation response, while high temperature activates the cell-wall integrity (CWI) response pathway (51–53). The three stresses related to the HOG pathway resulted in severe growth inhibition by LegA7 (Fig. 3A; compare Gal. + Sorb, NaCl or 20°C to Gal. in absence of stress), indicating growth inhibition by the effector is linked to activation of the HOG pathway. On the contrary, high temperature (37°C) resulted in suppression of the yeast growth inhibition by LegA7 at 30°C (Fig. 3A; compare 37°C to standard growth conditions). To further explore these results, we examined the yeast growth inhibition caused by LegA7, using yeast deletion mutants in the HOG pathway (*hog1* and *pbs2*) and CWI pathway (*mpk1* and *bck1*; Fig. 3B). Using standard yeast growth temperature (30°C, Glu) the four deletion mutants were indistinguishable from wild-type yeast. In contrast, at 37°C, the HOG pathway-related mutants (*mpk1* and *bck1*) still suppressed the LegA7-induced growth defect, whereas the two CWI pathway mutants (MPK1 and Bck1) failed to suppress this defect (Fig. 3C). These results indicate that yeast must activate the CWI pathway to suppress the growth defect mediated by LegA7 at 37°C, and in the absence of a component from this pathway, there is a growth defect at high temperatures.

**Figure 3.**
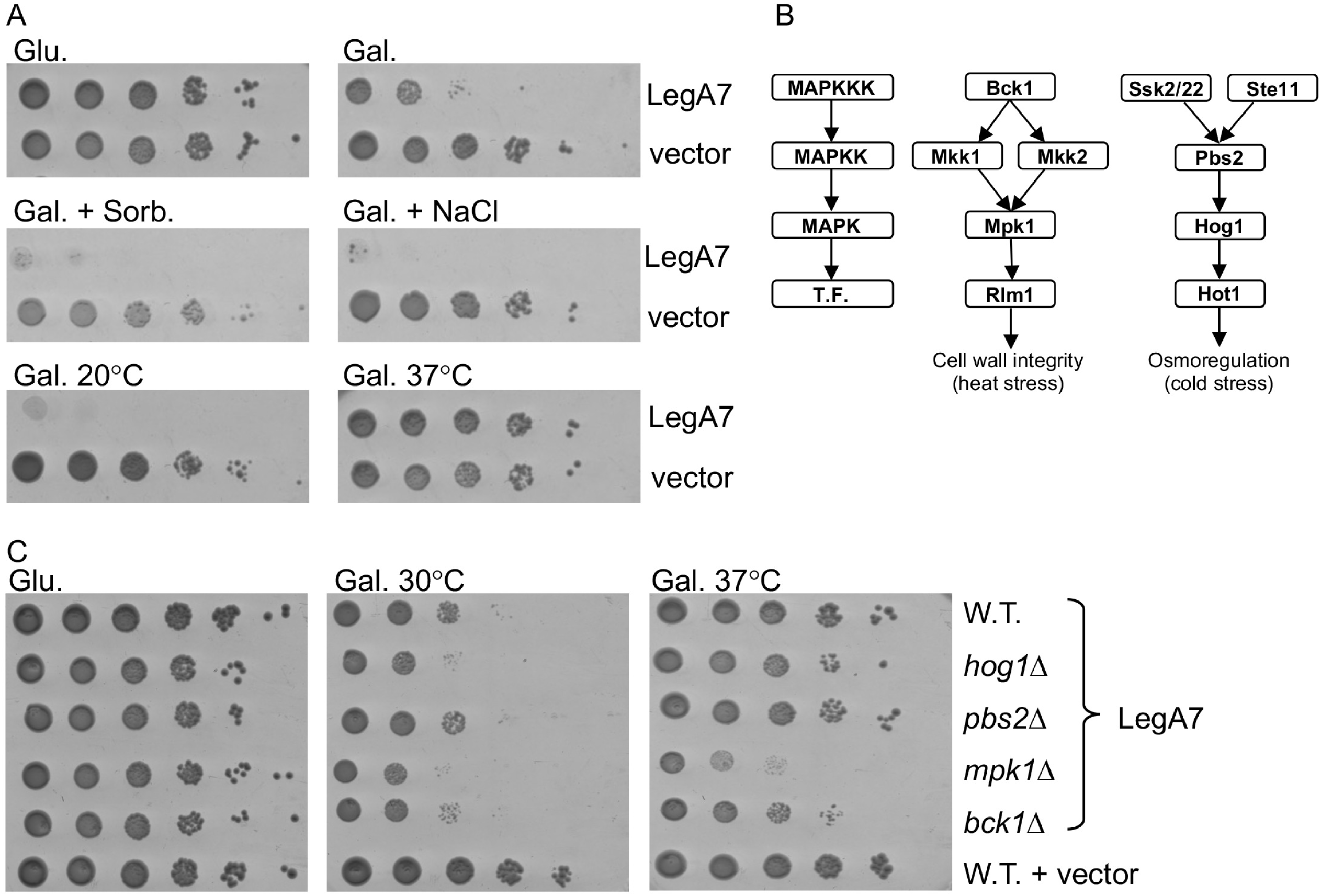
LegA7 affects MAPK pathways in yeast. **A.** The *L. pneumophila* LegA7 effector cloned under the GAL1 promoter was grown on plates containing glucose (Glu), galactose (Gal, inducing conditions), galactose supplemented with 1 M sorbitol (Gal. Sorb.), galactose supplemented with 0.7 M NaCl (Gal. NaCl) at 30°C and on SD plates containing galactose (Gal.) at 20°C and 37°C, in the wild-type *S. cerevisiae* BY4741. Ten-fold serial dilutions were performed, and the growth inhibition effect was compared to the one of the vector pGREG523 control (vector). **B.** Diagram of the yeast CWI MAPK pathway and the HOG MAPK pathway. The function of each protein is indicated on the left. T.F., transcription factor. C. Examination of the inhibition of yeast growth mediated by the LegL7 effector in deletion mutants of the CWI and HOG MAPK pathways. LegA7 was overexpressed in the wild-type *S. cerevisiae* BY4741 (W.T.) and the *hog1*, *pbs2*, *mpk1*, and *bck1* deletion mutants at 30°C and 37°C. Ten-fold serial dilutions were performed, and the growth inhibition effect was compared to the one of the vector pGREG523 control (vector).

### LegA7 shows sequence similarity to a family of bacterial cysteine protease effectors

Although the carboxyl-terminal of LegA7 has been shown to have a series of ankyrin-repeats, the amino-terminal has not been annotated, but was shown to harbor the LED010 domain present in other *Legionella* effectors (20). LED010 is one of a series of *Legionella* effector domains (LEDs) that was used to describe protein regions showing no similarity to known domains in database searches, but are conserved across multiple *Legionella* effector orthologous groups (20). In this analysis it was found that LegA7 was among a number of such effectors harboring the LED010 domain. We noted during BLAST searches that there was low similarity to HopN1, a *Pseudomonas syringae* type III secretion system translocated substrate that is similar to cysteine protease family members at key catalytic residues (54). The region of sequence similarity with HopN1 begins at the LegA7 Cys61 residue, which aligns with the predicted catalytic cysteine of the *Pseudomonas* protein (Fig. 4) (55). The cysteine in members of this family is part of a catalytic triad of essential amino acids that also includes a histidine and either a glutamate or an aspartate, usually located carboxyl-terminal to the cysteine (54). Scanning of the LegA7 sequence indicates that there are candidate residues that could be part of this catalytic triad and be required for the activity of this protein (Fig. 4).

**Figure 4.**
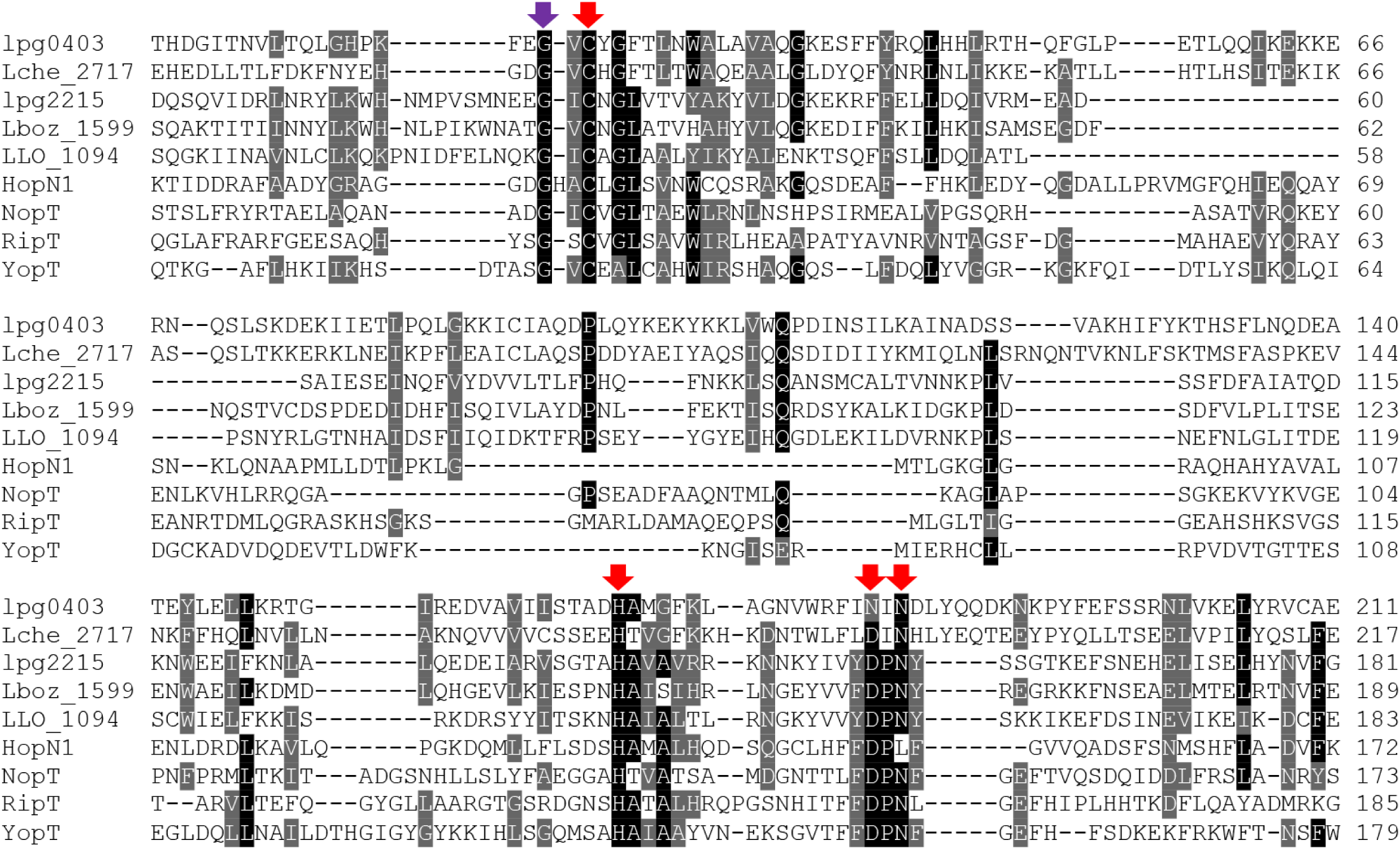
Similarity of LegA7 to bacterial cysteine protease family members. Sequence similarity alignment of LegA7 and several members of a bacterial cysteine protease family. Highlighted in red are the residues predicted to be part of the catalytic triad; highlighted in purple is the Glycine residue that came out in the mutagenesis screen (see text). Accession numbers are as follows: lpg0403 (LegA7) – *L. pneumophila,* AAU26500; Lche_2717 – *L. cherrii,* KTC80697; lpg2215 (LegA2) - *L. pneumophila*, AAU28280; Lboz_1599 – *L. bozemanae*, KTC74159; LLO_1094 – *L. longbeachae*, CBJ11449; HopN1 - *Pseudomonas syringae*, KPB86840; NopT - *Sinorhizobium fredii*, AAB91961; RipT - *Ralstonia solanacearum*, CBJ35895; and YopT - *Yersinia pestis*, WP_002213006.

After analysis of the LegA7 protein sequence, we selected residues C61, H205, N220, and N222 as candidate residues to make up this triad. Using site-directed mutagenesis, each residue was mutated to alanine. Mutations in all four residues described above were able to rescue the growth defect caused by LegA7 in yeast (Fig. 5A), with the efficiency of CFU formation for all strains being identical to the empty vector control (Fig. 5A,B). These mutants are consistent with C61, H205, N220, and N222 being important for LegA7 function and involved in a catalytic triad (Fig. 5C). The suppression of the defect was not a result of the mutations destabilizing the protein, because each of the mutant proteins showed higher steady-state levels of protein, than that observed for the wild-type protein, consistent with yeast tolerating these nontoxic proteins (Fig. 5B). In fact, the enhanced production of the mutants relative to wild-type is consistent with its loss of toxicity for yeast. This was also observed in mammalian cells, as the degradation of LegA7 that occurred after transfection of HEK293T (Fig. 2A) was greatly reduced if a Cys->Ala mutation was introduced at the predicted C61 residue (Supplemental Fig. S1), arguing for autodegradation.

**Figure 5.**
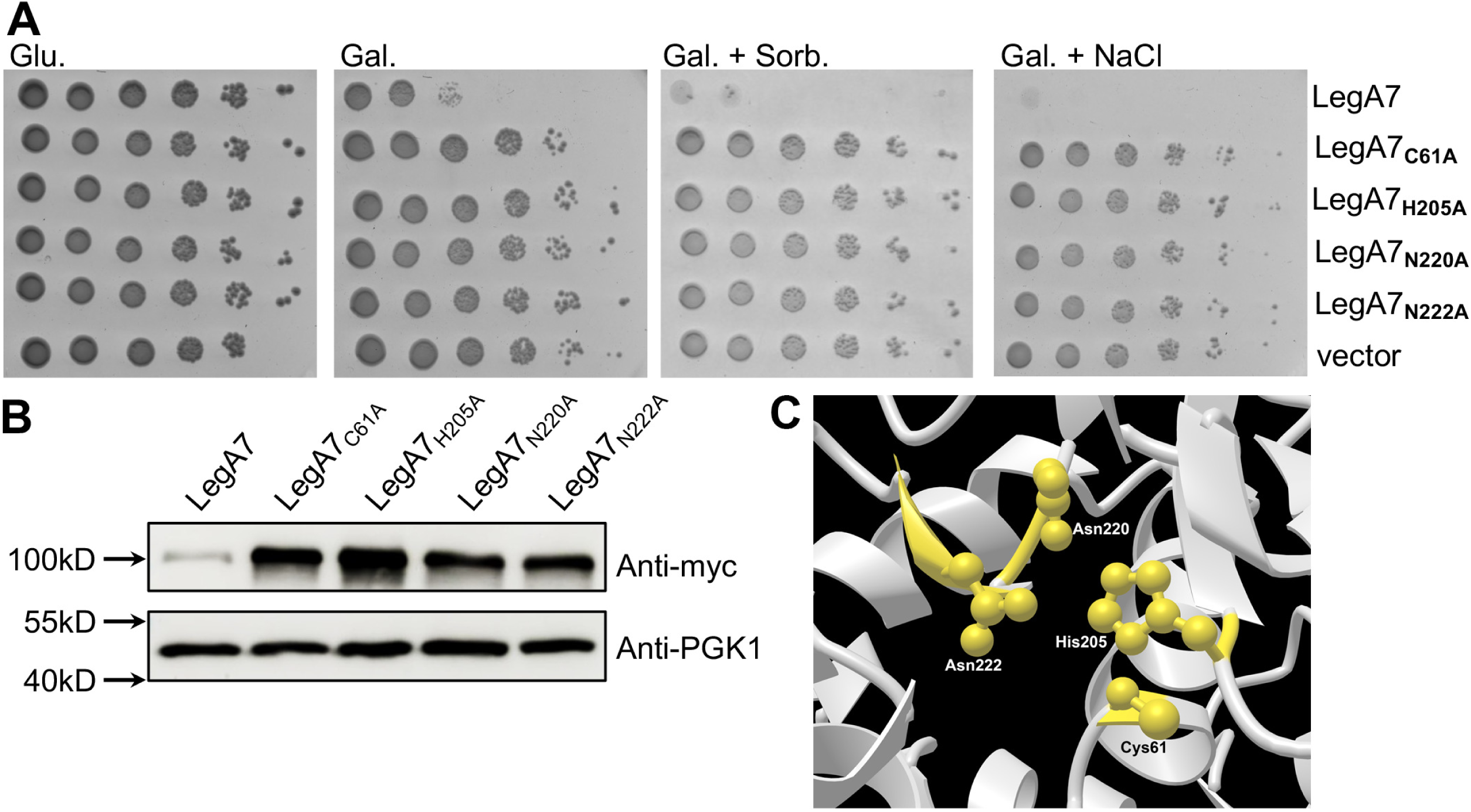
Genetic evidence for a catalytic triad in LegA7. **A.** Based on sequence similarity in Fig. 4, the residues C61, H206, N220, and N222 were selected as candidates for a catalytic triad. Point mutations were generated and plating efficiency on galactose (Gal.), galactose, and sorbitol (Gal. + Sorb.) galactose and NaCl (Gal. + NaCl) plates of yeast strains harboring the mutant derivatives was determined. Ten-fold serial dilutions were performed, and the growth inhibition effect was compared to the one of the vector pGREG523 control (vector). **B.** LegA7 point mutations do not reduce steady-state levels of protein. To induce gene expression in yeast, yeast strains were grown on SD plates containing galactose. Lysates were analyzed by immunoblot with antibodies against the *myc* epitope, using PGK1 for loading control.

### Mutagenesis screen to identify residues important for the LegA7 function

The carboxy-terminal of LegA7 is predicted to have five ankyrin repeats (residues 290-454) (49). Each ankyrin repeat contains two alpha-helices separated by beta turns, such that each 33-residue motif contains a beta turn-alpha helix-beta turn-alpha helix-beta turn. In addition to the five ankyrin repeats, there is an inter-domain region just amino-terminal to the ankyrin repeats. To determine if these repeats are important for LegA7 function, deletion mutations were constructed that lack the following: all five ankyrin repeats (Δ290-454); the carboxy-terminal four repeats (Δ331-454); the amino-terminal two repeats (Δ290-361); or the two amino-terminal repeats in conjunction with the inter-domain region (Δ264-361) (Fig. 6). None of the deletions caused a marked growth defect when introduced into yeast and grown on galactose, galactose and sorbitol, or galactose and NaCl plates (Fig. 6A). The lack of growth defect did not appear to be a result of the LegA7 derivatives being unstable because only the derivative lacking all five ankyrin repeats showed low steady-state levels of protein (Fig. 6C).

**Figure 6.**
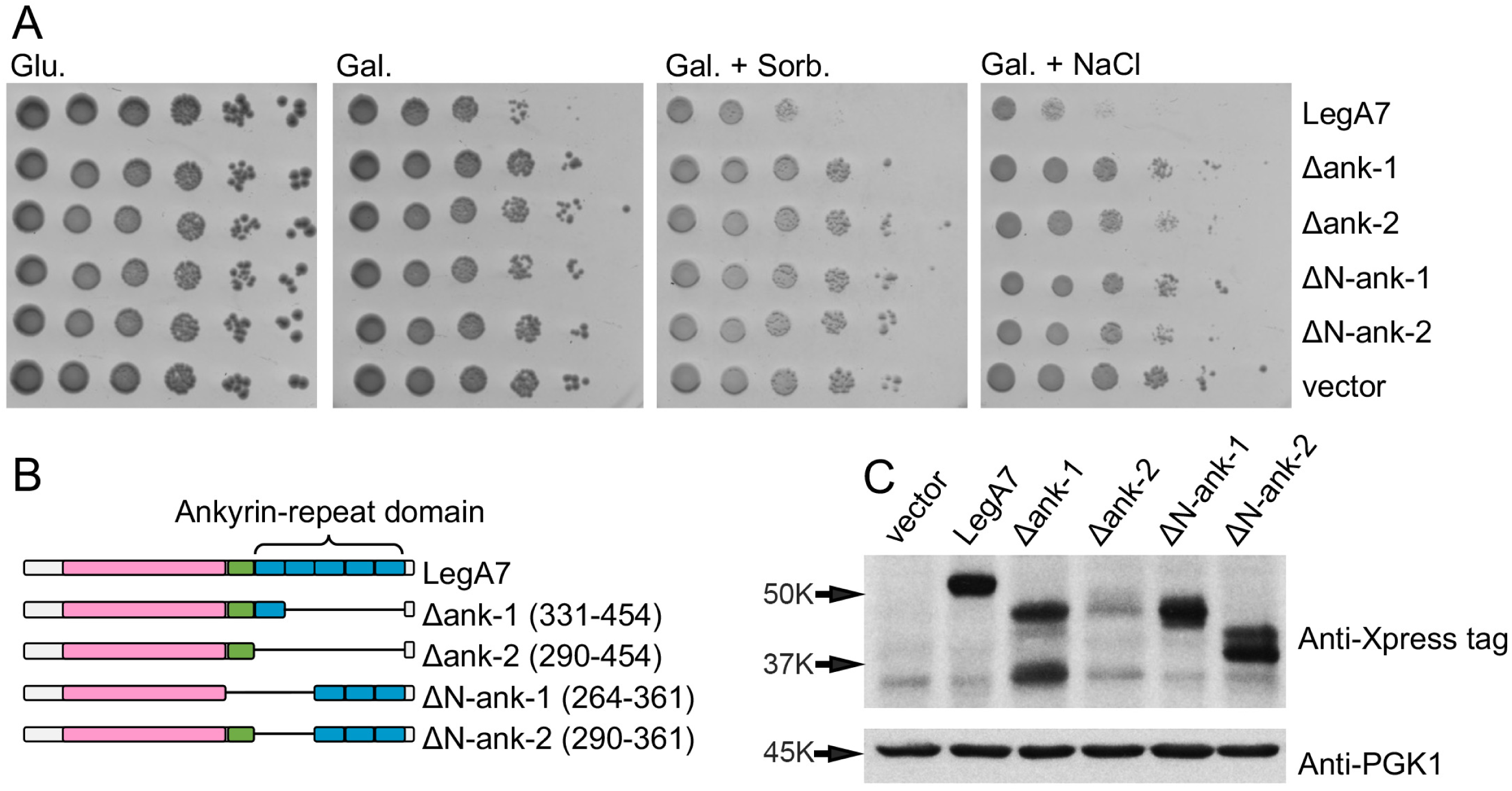
Ankyrin repeats are required for yeast growth inhibition. **A.** Deletion of the ankyrin repeats relieves LegA7-induced yeast growth inhibition. LegA7 (502 amino acids long) is predicted to have five ankyrin repeats from residues 290-454 and an inter-domain region upstream. To demonstrate that the ankyrin repeat domain is important for LegA7 function, the entire ankyrin repeat domain (residues 290-454), the four carboxy ankyrin repeats (residues 331-454), the first two amino-terminal ankyrin repeats (residues 290-361), or the first two amino-terminal ankyrin repeats and the inter-domain region (residues 264-361), were deleted from LegA7 and plating efficiency on galactose (Gal.), galactose and sorbitol (Gal. + Sorb.) galactose and NaCl (Gal. + NaCl) plates of yeast strains harboring the mutant derivatives was determined. **B.** Schematic of *legA7* showing the cysteine peptidase domains (pink box), the inter-domain (ID) region (green box), and the ankyrin repeat domain-containing five repeats (blue boxes). **C.** Western blot demonstrating expression of LegA7 and LegA7 deletions. To induce gene expression in yeast, overnight cultures were back-diluted into a medium containing 2% galactose for 5 hours. Lysates were analyzed by immunoblot with antibodies against the Xpress epitope and PGK1 for loading control. The ΔAnk (331–454) protein (partial ankyrin-repeat domain deletion) is predicted to migrate around 43 kDa. The ΔAnk (290–454) protein (full ankyrin-repeat domain deletion) is predicted to migrate around 39 kDa.

To further characterize residues important for LegA7 function, we screened for mutations on a LegA7 expression plasmid that no longer inhibited yeast growth, using a strategy that prevented the isolation of premature termination mutants. To this end, we constructed a plasmid with the *HIS3* gene (histidine biosynthesis) placed at the 3’-end of *legA7*, and introduced it into a *his3* auxotrophic strain. We then selected for mutations on the plasmid that lost the ability to interfere with yeast growth, but retained the ability to grow in the absence of histidine (56). This strategy selected against frameshifts, stop codons, or GAL1 promoter mutations, each of which would result in enhanced growth relative to strains harboring the *legA7* gene (Fig. 7A). Using a similar strategy to isolate IDTS point mutations, the absence of the HIS3 co-selection resulted in the vast majority of mutants being noninformative truncations, strongly supporting the use of this approach (57).

**Figure 7.**
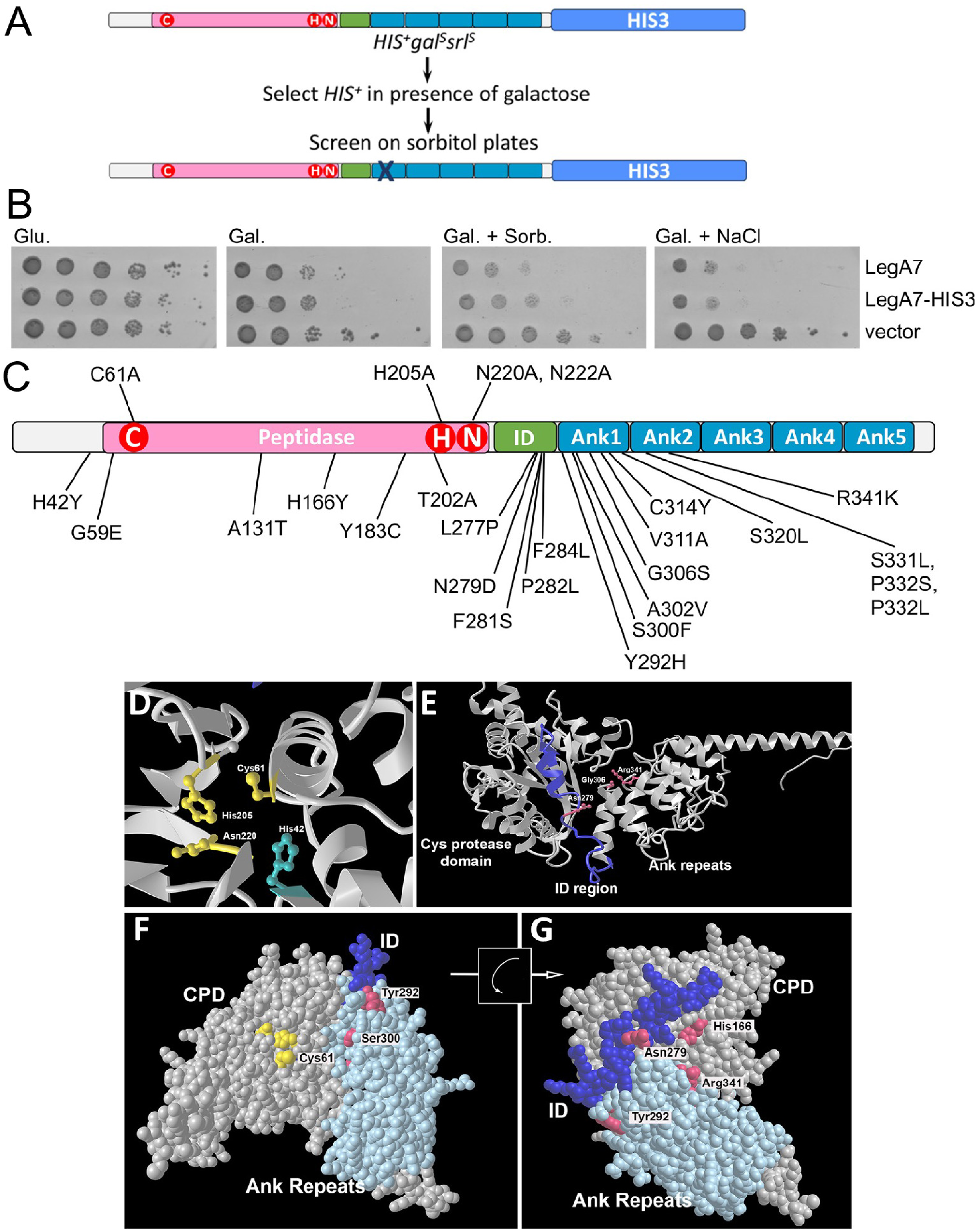
Identification of residues in the amino-terminal ankyrin repeats that are essential for exerting yeast growth inhibition. **A.** Selection and screening for LegA7 mutants that fail to cause hyperosmotic stress. Plasmid harboring LegA7-HIS3 protein fusion was mutagenized (Materials and Methods), transformed into yeast and selected for HIS3^+^ in presence of galactose. Surviving colonies were streaked onto sorbitol-containing plates on histidine dropout medium. Gal: galactose; srl: sorbitol. **B.** Demonstration that *legA7-HIS3* causes growth inhibition when expressed in yeast. The panel shows the plate assay of the growth of LegA7-HIS3 fusion as it compares to LegA7 alone. The *L. pneumophila* LegA7 and LegA7::HIS3 were cloned under the GAL1 promoter were grown on plates containing glucose (Glu), galactose (Gal, inducing conditions), galactose supplemented with 1 M sorbitol (Gal. Sorb.), galactose supplemented with 0.7 M NaCl (Gal. NaCl) at 30°C. Ten-fold serial dilutions were performed, and the growth inhibition effect was compared to the one of the vector pGREG523 control (vector). **C.** Clustering of mutations in the amino-terminal region of the ankyrin repeats that relieve yeast growth inhibition. Schematic of LegA7 showing the cysteine peptidase domains (pink box), the inter-domain (ID) region (green box), and the ankyrin repeat domain-containing five repeats (blue boxes). The residues marked with red circles are the residues that make up the putative catalytic triad. The mutations indicated below schematic were identified in the *legA7-HIS3* mutagenesis. The mutations indicated above are the directed point mutations that were used to identify the putative triad in Fig. 5. It is likely that Genbank incorrectly annotated the translation start site as 13 codons upstream from the designated start site in this panel. The transcriptional start site is downstream from the Genbank annotation, so the residue numbering system in this panel begins with the first start codon available after the transcription start, where there is also a consensus ribosome binding site. **D.** Predicted LegA7 catalytic triad (gold), including site of His42 residue (blue) mutation isolated in selection strategy. E. The ID region links catalytic domain to Ank repeats. Noted are surface-exposed residues that were mutation sites isolated in panel A. F. Space-filling model of LegA7 residues 1-465 showing arrangement of Ank repeats and catalytic site. G. Image of panel F, rotated as noted, showing back surface relative to catalytic domain, with sites of mutations described in panel C.

A plasmid encoding the *legA7-HIS3* protein fusion was subjected to mutagenesis by passage within an *E. coli* mutator strain (Fig. 7A; Materials and Methods). Mutated plasmids were then introduced into yeast, selecting on medium containing galactose inducer and lacking histidine, to allow selection for intact *legA7-HIS3* protein fusions with increased viability after induction of expression (Fig. 7B). After three days incubation, the wild-type *legA7-HIS3* fusion showed little growth, so colonies were retained from two mutagenized pools at this timepoint. Of these, 36 were shown to have strong growth on sorbitol-containing high-osmolarity medium and retained a functional *legA7-HIS3* protein fusion (Table 2). The high osmolarity resistant mutants fell into two categories: a few were found in the region surrounding the catalytic triad (G59A, T202A), but the majority were found in the inter-domain region and the two most amino-terminal ankyrin repeats (Fig. 7C). Of the residues near the catalytic triad, the most notable was G59. Although not part of the triad, the Gly residue is highly conserved in members of the bacterial cysteine protease family, located 2 or 3 residues upstream from the catalytic cysteine (Fig. 4, purple arrow). Two additional abundant sites for mutant isolation were H166 and S300, which were hit seven and six times, respectively. There were 17 additional mutations in the two amino-terminal ankyrin repeats and five mutations in the inter-domain region. These 22 mutations emphasize the importance of the inter-domain region and the amino-terminal region of the ankyrin repeats for LegA7 function. In contrast, no mutations selected in this fashion were found in the three carboxy-terminal ankyrin repeats.

**TABLE 2.** LegA7 mutations that allow enhanced survival on high osmolarity medium.

| Nucleotide change <sup>a</sup> | AA change | Region altered <sup>b</sup> | Times isolated | Relative growth <sup>c</sup> |
| --- | --- | --- | --- | --- |
| C124T | H42Y | Amino terminal | 1 | +++ |
| G176A | G59E | Amino terminal | 1 | +++ |
| G391A | A131T | Amino terminal | 1 | +++ |
| A398T; G904A | D133V; A302T | Amino terminal | 1 | +++ |
| C496T | H166Y | Amino terminal | 7 | +++ |
| A548G | Y183C | Amino terminal | 1 | +++ |
| A604G | T202A | Amino terminal | 1 | ++ |
| T831C | L277P | ID region | 1 | ++ |
| A835G | N279D | ID region | 1 | ++ |
| T842C | F281S | ID region | 1 | ++ |
| C845T | P282L | ID region | 1 | ++ |
| T850C | F284L | ID region | 1 | ++ |
| T874C | Y292H | Ank repeat 1 | 1 | +++ |
| C899T | S300F | Ank repeat 1 | 6 | +++ |
| C905T | A302V | Ank repeat 1 | 1 | +++ |
| G916A | G306S | Ank repeat 1 | 1 | +++ |
| T932C | V311A | Ank repeat 1 | 1 | +++ |
| A936G; C994T | K312K (silent); P332S | Ank repeat 1 | 1 | +++ |
| G941A | C314Y | Ank repeat 1 | 1 | ++ |
| C959T | S320L | Ank repeat 2 | 2 | +++ |
| C992T | S331L | Ank repeat 2 | 2 | +++ |
| C995T | P332L | Ank repeat 2 | 1 | +++ |
| G1022A | R341K | Ank repeat 2 | 1 | +++ |

<sup>a</sup> Listed are mutations isolated by selecting for enhanced growth on sorbitol-containing medium of *legA7-HIS3* fusion strains.
<sup>b</sup> ID – Inter domain region, and Ank – Ankyrin repeat.
<sup>c</sup> Relative degree of growth is noted by +++, which is indistinguishable from the C61A mutation.

To evaluate the putative active site mutations and determine if any of the mutations in the inter-domain and ankyrin repeat regions could directly interface with the target of this activity, the residues altered were modeled with AlphaFold 2.0, using the ColabFold program (Materials and Methods; (58, 59)). The predicted 3-dimensional structure is consistent with C61/H202/N220 forming a catalytic triad (Fig. 4 and Fig. 7D), with the catalytic Cys residue in close apposition to His205. In addition, the defective H42Y mutation is located in a residue that interfaces with Glu58 and Gly59 directly abutting the active site (Fig. S2). The Tyr substitution is situated so that it could impinge on the active site, preventing substrate access or altering interactions between catalytic residues.

The mutations located in the inter-domain (ID) and ankyrin repeat (Ank) regions are consistent with LegA7 activity being regulated by an interaction surface that is distant from the catalytic triad. Many of these mutations alter hydrophobic or small sidechain amino acids that appear poorly accessible to water, likely causing local structural alterations that block presentation of a binding interface to host proteins. Among the few mutations in clearly surface-exposed residues, three were aligned with each other on each side of a cleft, with Ank1/2 on one side, and the catalytic domain/ID region on the other side (Fig. 7E). The clustering of these mutations (Asn279, Gly300, Arg341) is consistent with the formation of a binding surface for eukaryotic substrates, with the Asn and Arg residues participating in interprotein associations. This putative binding cleft is on the opposite side of the protein from the catalytic site, indicating that this surface may not be involved in target association with the catalytic site (Figs. 7F,G; note flip). Rather, this binding surface is oriented similarly to the interface between the *L. pneumophila* VipD phospholipase and the host Rab5 activating protein (60), consistent with host protein binding to the ID and Ank1/2 regions resulting in activation of LegA7 catalysis.

## DISCUSSION

*Legionella pneumophila* has evolved an arsenal of methods to manipulate the host cell to survive and replicate intracellularly. To this end, *L. pneumophila* translocates hundreds of IDTS proteins into the host cell through the Icm/Dot type IVb secretion system. Among these proteins are ones known to activate NFκB, such as LnaB, and the kinase LegK1 (34, 61), as well as at least five protein synthesis inhibitors that activate both NFκB and MAPKs (28, 62). To identify other IDTS that alter the host cell stress response, we introduced a 259-member plasmid bank of *L. pneumophila* IDTS into mammalian cells and screened for proteins that caused phosphorylation of the stress-activated MAPK SAPK/JNK (63). A complimentary screen was performed in yeast, by overexpressing IDTS proteins in yeast in the absence or presence of sorbitol, to identify IDTS proteins that amplify defects on a medium that activates a stress pathway requiring MAPK activity (53). Both screens led us to focus on a single IDTS, LegA7, which seems to impinge on a stress response pathway across evolutionarily diverse hosts.

The results of the yeast growth inhibition by LegA7 were complex and found to be dependent on growth conditions. Induced expression of LegA7 resulted in moderate growth inhibition of yeast on solid laboratory growth medium without additives and at standard temperatures (30°C; Fig. 3A). Growth inhibition was greatly exacerbated by the addition of osmolytes such as sorbitol or NaCl but was completely suppressed by incubation on standard medium at high temperature (37°C; Fig. 3A). Our hypothesis is that LegA7 inhibits the activity of components in both the Hog1 and Mpk1 pathways (Fig. 3B). On addition of an osmolyte, the HOG pathway is activated to ensure survival, but LegA7 presumably interferes with HOG pathway activation, resulting in growth inhibition. Incubation at 37°C results in transcriptional activation of the MAP kinase pathway, possibly leading to a titration effect that allows the accumulation of Hog1 and Mpk1 pathway substrates that are left untargeted by LegA7. This hypothesis is strongly supported by the fact that at high temperature, no rescue of yeast growth was observed in either the *mpk1* or *bck1* deletion mutants. Presumably, these mutations allow the Hog1 pathway to be targeted more efficiently at 37°C by LegA7 compared to WT strains, as there is a reduction in the substrate concentration, with consequent saturating targeting of this pathway.

LegA7 was previously identified as a substrate of the Icm/Dot secretion system due to its ankyrin repeats (64) and it is homologous to a cysteine protease domain at its amino-terminal (Fig. 4). Mutagenesis of conserved residues of this cysteine protease domain (Fig. 5) supports the model that these residues form a catalytic triad similar to that found in members of this family (54). Cysteine protease domain family members are associated with a variety of catalytic functions in addition to performing proteolysis, such as small molecule transferases (54, 65). Most of these activities are unknown and await identification. One of the most important features of this family is that members show high substrate specificity, and target single residues in their substrates. This is exemplified by the defining members, *Pseudomonas syringae* AvrPphB and *Yersinia* YopT, which are type III secretion system translocated substrates (55). AvrPphB is a protein with a papain fold that has a protease activity targeting a plant protein associated with innate immune signaling (66, 67). YopT cleaves membrane-bound Rho GTPases just upstream from their acylation sites, resulting in the release of these proteins from the plasma membrane and disruption of the host actin cytoskeleton (55).

Cysteine protease domains like that identified in LegA7 are found in another *L. pneumophila* effector (LegA2-Lpg2215) and in putative effectors in other *Legionella* species (Fig. 8A). Each of these proteins (LegA7, LegA2, and the other IDTS proteins presented), belong to an orthologous group, all harboring the cysteine protease domain at the amino-terminal part of the protein, with ankyrin repeats of varying numbers found at the carboxyl terminal (Fig. 8A). In addition, one of these IDTS proteins also harbors a predicted phosphatidylinositol 3-phosphate (PI3P) binding domain (LED027), which was previously shown to bind PI3P in other *L. pneumophila* effectors (68), possibly directing effectors to the LCV surface (69–71). The catalytic triads identified in LegA7 and the putative IDTS proteins presented are very similar to the ones found in numerous type III secreted effectors (Fig. 8B). However, in LegA7 an asparagine residue is located in the position that is usually occupied by an aspartic acid residue in the catalytic triad in the type III effectors (Fig. 8B). The asparagine residue that is critical for the function of LegA7 has recently been shown to be critical for other unrelated cysteine proteases that also harbor conserved cysteine and histidine residues in their catalytic triads (72). At this point, there is no clear sequence motif that distinguishes protease from transferase activity, so the presence of the asparagine in LegA7 should not be considered diagnostic of a particular activity. For instance, the *Yersinia* YopJ type III effector has the typical catalytic triad of a cysteine peptidase, but it functions as an acetyltransferase that targets MAPK kinases, preventing their activation (73).

**Figure. 8.**
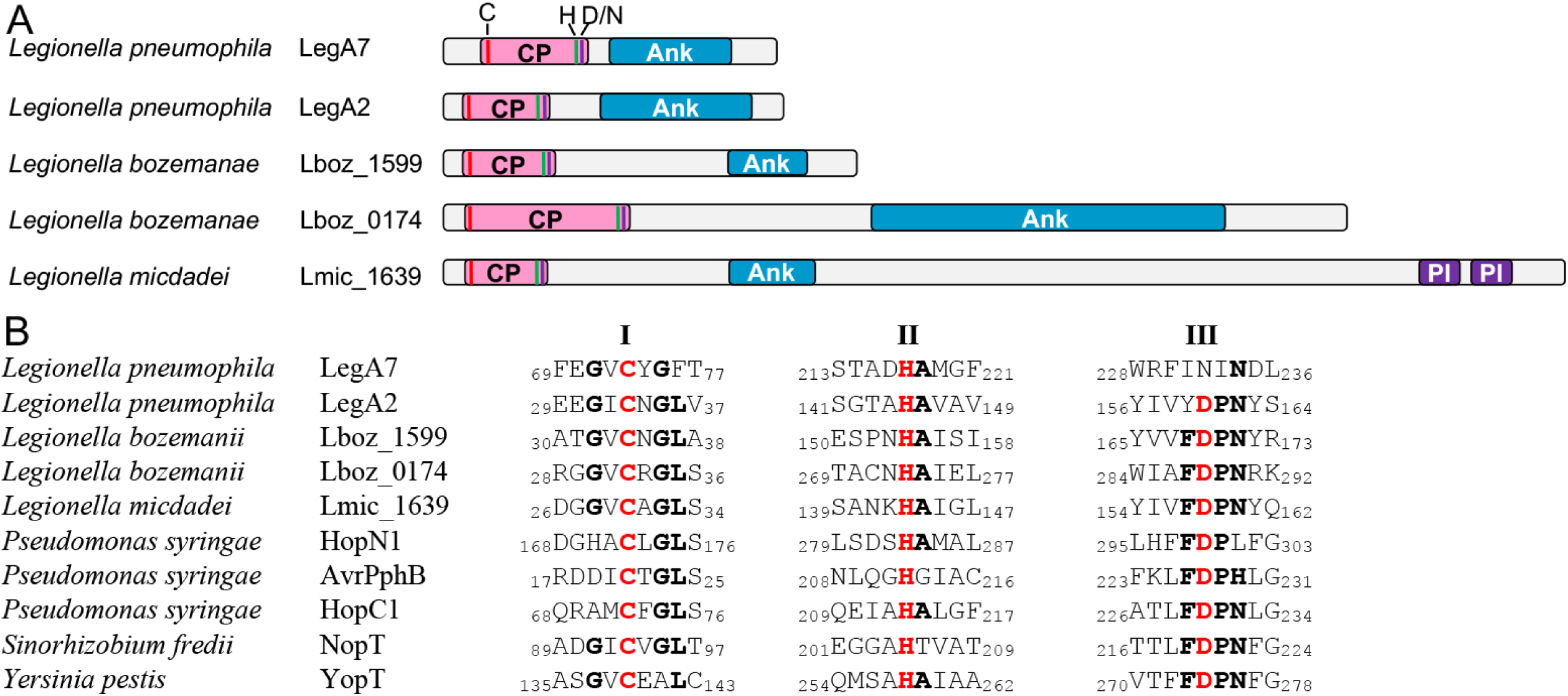
The cysteine protease domain of Icm/Dot and type III effectors. A. Domain architecture of *Legionella* IDTS proteins harboring the cysteine protease (CP) domain. The known domains of each IDTS that contain the cysteine protease domain are shown. The domains presented are Ank - ankyrin domain and PI - PI3P-binding domain. B. The catalytic triad of the cysteine protease domain. Amino acid sequence alignments of cysteine protease catalytic triad I to III of five IDTS and predicted IDTS proteins as well as type III secreted effectors. The numbers indicated the position of the amino acids present in each of the proteins. Accession numbers are as follows: LegA7, AAU26500; LegA2, AAU28280; Lboz_1599, KTC74159; Lboz_0174, KTC77221; HopN1, PB86840; AvrPphB, SPD82586; HopC1, AAO54131; NopT, AAB91961; and YopT - WP_002213006.

A mutagenesis screen to identify residues important for LegA7 function provided additional information about other domains important for the function of LegA7 (Fig. 7C). Sequence similarity alignments with other cysteine protease domains predicted a catalytic triad (Fig. 4) and this prediction was verified by mutations in the predicted catalytic residues (Fig. 7B; Table 2). Furthermore, tertiary structure prediction from AlphaFold 2 indicated that the C61/H205/N220 triad formed a compact pocket in the catalytic domain, consistent with the mutant analysis (Fig 7D). Notably, a His42➔Tyr mutation in the catalytic domain was isolated by selection in yeast, at a site not predicted to be involved in catalysis. The structural model, however, predicted that this residue was at the base of this catalytic pocket (Fig 7D). When the insertion of the Tyr at residue 42 was modeled compared to the WT His residue, it was found to impinge on neighboring residues Glu58 and Gly59 in the pocket, perhaps distorting the site or blocking access to the catalytic residues (Fig. S2), thus providing a molecular explanation for the isolation of this mutation.

We isolated five mutations in the inter-domain region and 11 mutations in the two ankyrin repeat motifs immediately downstream from this region that reduced yeast growth inhibition. Although many of the mutations in these two regions appeared to be in buried residues or possibly altering structure, there was a series of residues predicted to be surface exposed on a face of the protein where the inter-domain (ID) region comes in contact with the Ank1 and Ank2 repeats (Fig. 7E). The region altered by these mutations is predicted to be turned 180° away from the catalytic pocket, making it unlikely that the substrates of the protease domain were binding on this face of the protein. It is more likely that this region allows LegA7 to target to a cellular locale where it can access its substrates. Alternatively, binding to this region by a host or bacterial protein could activate LegA7, allowing either localization- or time-dependent activation of the catalytic triad. *Legionella* translocated proteins are known to be highly regulated, both by other translocated proteins called metaeffectors (74), or by host proteins. The fact that this proposed regulatory surface appears to be on the face opposite from the catalytic triad (Fig. 7F,G) is not unusual, and has been previously observed in the crystal structure of the *L. pneumophila* VipD patatin family phospholipase, which is activated by Rab5 (60).

One of the puzzles of the point mutation screen is that no lesions were identified in the carboxyl- terminal three ankyrin repeat domains (Table 2; Fig. 7C). Some insight into this result may be given by the crystal structure of AnkX, an *L. pneumophila* IDTS ankyrin repeat-containing protein having phosphocholine transferase activity (39, 75, 76). Based on primary sequence information, AnkX is predicted to have up to 12 ankyrin repeats arrayed downstream from the catalytic domain, similar to the domain arrangement of LegA7. The crystal structure of the phosphocholine transferase domain indicates that the amino-terminal four ankyrin repeats are involved in intramolecular interactions that support the catalytic activity of AnkX (76). A proteolytic cleavage product that retains only the amino-terminal four repeats and the phosphocholine transferase region retains full activity and substrate specificity, consistent with the carboxyl-terminal repeats being dispensable for activity (76). Presumably, the ankyrin repeats in this protein are divided into an amino-terminal region involved in intramolecular interactions, with the carboxyl-terminal region providing intermolecular interactions that contribute to the localization or targeting of protein. By analogy with AnkX, the two amino-terminal repeats of LegA7 may be involved in intramolecular interaction supporting the activity of the protein, with the three carboxyl-terminal repeats involved in spatial targeting of the protein or processes unrelated to lethality in yeast.

The physiological role and substrates of LegA7 are unclear, because the activation of JNK (Fig. 2) could indicate direct targeting of this pathway or a regulatory response of host cells to stress induced by the *Legionella* protein. Previous work on *L. pneumophila* argues for the latter model, as the Lgt1 and Lgt3 translation elongation inhibitors are among the most robust *Legionella* inducers of the MAP kinase response (28, 62). We favor a model in which LegA7 has a regulated protease activity because the protein appears to undergo Cys61-dependent autodegradation after introduction into eukaryotic cells (Supplemental Fig. S1) and a region of the protein that is distant from the catalytic triad is required for the biological effects observed in this work (Fig. 7). This model requires that the host collaborate with the pathogen, relieving negative regulation of the protein, to drive cleavage of a critical substrate that supports microbial growth and results in host cell stress.

We have determined that LegA7 appears to activate at least one host cell pathway that, when disrupted, results in yeast growth inhibition. Using a genetic strategy, we were able to obtain evidence that yeast growth inhibition likely results from an enzymatic activity at the amino-terminal end of this protein that is modulated by a subset of ankyrin repeats. Future work will be required to identify the substrate of LegA7 activity and enumerate the pathways that are misregulated by the targeting of this substrate.

## MATERIALS AND METHODS

### Strains and media

Yeast and bacterial strains, plasmids and primers used in this study are listed in Datasets S2, S3 and S4, respectively. For *E. coli* strains, ampicillin was added to 100 µg·ml^-1^, kanamycin was added to 30 µg·ml^-1^. Yeast strains were grown in synthetic defined (SD) dropout medium supplemented with 2% glucose, or galactose as indicated in the text.

### Screen for MAPK activation in mammalian cells

Plasmids containing Icm/Dot translocated substrate genes fused to *gfp* were constructed by inserting fragments into pDONR221 (Invitrogen) and transferring the inserts into pDEST53 (pCMV-GFP) by using the Gateway™ system, as we previously described (34). The inserts in the original pDONR211 constructions were sequenced, and the appropriate recombinants were transferred into the GFP-expressing plasmid (34). The GFP fusion constructions, in which GFP-IDTS fusions were under the control of the CMV promoter, were analyzed by restriction digestion, the insertions were sequenced, and the plasmids were purified using Ultra-Pure Miniprep kits (Qiagen) for use in transfections.

To screen for MAPK activation, HEK293T cells were seeded at a density of 1 x 10^6^ cells per well of 12-well dishes and left overnight to adhere. Cells were transfected with 500 ng of each plasmid using 0.4 µl of Fugene 6 according to the manufacturer’s instructions (Promega) for 40 hours. Each well was washed, solubilized in SDS sample buffer (2% SDS, 50 mM TRIS-HCl (pH = 6.8), 0.1% Bromphenol Blue, 10% glycerol), boiled for 2 min., loaded onto two SDS-PAGE gels, and transferred to filters for immunoprobing. One filter was used to probe for relative expression levels of each of the fusion constructions, using anti-GFP polyclonal serum A-11122 (Invitrogen). The other filter was immunoprobed using anti phospho-ERK antibody 9101 (Cell Signaling), anti phospho-JNK antibody 4668 (Cell Signaling), or anti phopho-p38 antibody 4631 (Cell Signaling). Filters were scanned by densitometry and the relative phosphorylation level of each MAPK member was determined relative to anti α-tubulin antibody T9026 (Sigma) loading control. Images were inverted and quantified using Adobe Photoshop.

Analysis of MAPK phosphorylation samples resulted in a few transfections that showed large amounts of JNK phosphorylation relative to the control empty vector, giving the overall dataset a large standard deviation. Therefore, to expand the number of *L. pneumophila* candidates that cause alterations in JNK phosphorylation relative to empty vector control, the median and the median absolute deviation (MAD) of each sample tested were determined (77). A MAD score was then determined for each sample as (X_i_ – median)/MAD, in which X_i_ = amount of phosphorylation relative to the control of a particular sample. Samples that had MAD scores ≥ 3.5 were considered to be expressing candidate *L. pneumophila* proteins that cause enhanced activation of JNK.

### Quantification of MAPK activation in mammalian cells

For quantitative analysis of MAPK activation in mammalian cells, HEK293T cells were seeded at a density of 1 x 10^6^ cells per well of 12-well dishes and allowed to adhere overnight. Cells were transfected with 500 ng of each plasmid using 1.5 µl of Fugene HD according to manufacturer’s instructions (Promega) for 40 hours. Each well was washed, solubilized in 2x SDS sample buffer (125 mM Tris-HCl (pH = 6.8), 20% glycerol, 4% SDS, 2% 2-ME, 0.001% bromophenol blue), boiled for 5 minutes, loaded onto three SDS-PAGE gels, and transferred to PVDF membranes for immunoprobing. One filter was used to probe for relative expression levels of each of the fusion constructions, using anti-GFP polyclonal serum A-11122 (Invitrogen). The second filter was immunoprobed with anti phospho-JNK antibody 4668 (Cell Signaling). The third filter was immunoprobed with anti α-tubulin antibody T9026 (Sigma). The appropriate HRP-conjugated secondary antibodies (Invitrogen) were used for ECL detection. The filters were exposed to film and images of the films were taken using Kodak Image Station.

To determine the normalized signal intensity, the signal intensities of phospho-SAPK/JNK for *legA7*, lpg0030, lpg0059 and empty vector were normalized to the intensity of the loading control alpha-tubulin for each particular sample. The average and standard error were calculated for each strain. To determine ‘Relative Expression’ over the empty vector, the average normalized signal intensity of each strain was divided by the average normalized empty vector signal intensity. Data are the mean of three samples ± S.E.

### Yeast growth assays

*L. pneumophila* effector-encoding genes were cloned under the GAL1 promoter in the pGERG523 yeast overexpression vector. Plasmids were transformed into yeast cells using a standard lithium acetate protocol (78), and transformants were selected for histidine prototrophy on minimal SD dropout plates. Resulting transformants were then grown overnight in liquid SD culture medium at 30°C, cell number was adjusted, and a series of 10-fold dilutions were made. The cultures were then spotted onto the respective SD dropout plates containing 2% glucose or galactose. When indicated, the plates were supplemented with 0.7 M NaCl (Merck) or 1 M sorbitol (Sigma).

### Construction of 13 **x** myc fusions

The pGREG523 vector was used for the overexpression of 13 x *myc*-tagged effectors in yeast (79). This vector contains a polylinker under the yeast GAL1 promoter at the end of a 13 x *myc* tag.The *L. pneumophila* genes examined were amplified by PCR using a pair of primers containing suitable restriction sites (Dataset S2). The PCR products were subsequently digested with the relevant enzymes and cloned into pGREG523 to generate the plasmids listed in Dataset S3. The plasmid inserts were sequenced to verify that no mutations were introduced during the PCR.

### Site-directed mutagenesis and construction of deletion mutants

Site-specific mutants in the putative peptidase domain of *legA7* were constructed by the PCR overlap-extension approach on the *legA7* gene inserted into pGREG523 (80) as previously described, using the primers listed in Dataset S2.

Deletions of LegA7 ankyrin repeats were performed, using oligonucleotides overlapping deletion junctions. Primers containing the desired deletions and complement were used to amplify the entire plasmid sequence using *Pfu*Ultra II fusion HS DNA polymerase (Agilent). After PCR, the product was *Dpn*I treated to digest the parental DNA template. 5µL of the *Dpn*I-digested PCR reaction was then transformed into Ca_2_Cl-competent DH5alpha *E. coli*, allowing the linearized DNA to be recircularized by the *E. coli* cells (81). Mutants constructed were confirmed by Sanger sequencing (GeneWiz Azenta, South Plainfield, NJ). Plasmids harboring deletions mutants described in Fig.6 and Supplemental Fig. S3, were sequenced entirely (Plasmidsaurus, Eugene, OR), with sequences and plasmids being deposited with AddGene (ID numbers: 216518-216522).

### Western blot analysis

For all protein fusions examined in yeast, the formation of a fusion protein with a proper size was validated by Western blotting using anti-*myc* antibody 9E10 (Santa Cruz Biotechnology), and anti-Xpress tag antibody R91025 (Invitrogen). Anti-PGK1 antibody 22C5D8 (Invitrogen) was used as a loading control.

### Mutagenesis screen

To isolate random mutations in the *legA7* gene, the *HIS3* gene was placed directly downstream and in-frame with the *legA7* gene (removing the cognate stop codon) in the plasmid pYES2/NTA (URA selection) using homologous recombination to generate a *legA7*-*HIS3* fusion. The BY4741 strain transformed with the *legA7-HIS3* fusion pYES2/NTA was verified to show a stronger growth defect than the plasmid harboring *legA7* alone, as demonstrated by spotting assay. Mutagenesis was performed by transforming the plasmid into the *E. coli* XL1-Red mutator strain (*mutS mutT mutD*) (see Dataset S4). After growth in culture for 16 hours, DNA was isolated from two independent cultures and transformed into *S. cerevisiae* BY4741, selecting for growth on SD (galactose)-URA-HIS drop-out medium. Strains that gave higher viability or larger colony size than the parental *legA7-HIS3* protein fusion-containing strain on sorbitol-containing medium were retained for further analysis.

### Modeling of mutation sites

The full-length LegA7 was submitted to the ColabFold program (59) to allow analysis by AlphaFold 2.0 (58). The five ranked models that were returned provided similar results, so the Rank 1 model was used for further analysis. The PDB file generated (Dataset 4) was displayed in iCn3D (82), allowing sites of mutations to be identified and relative orientation of domains to be evaluated.

## ACKNOWLEDGMENTS

EJC was partially supported by MSTP training grant T32GM008448. AD and AE were supported by NIAID fellowships F32-AI084202 and F32AI074193 respectively. This work was supported in part by grant 2019101 from the United States–Israel Binational Science Foundation (RI and GS). The Work was supported by Howard Hughes Medical Institute and NIAID award R01AI46245. We thank Dr. Matt Heidtman for help during the early stages of this work, and Drs. Edward Geisinger, Dervla Isaac, Tamara O’Connor, and Kim Davis for review of the manuscript.

## REFERENCES

1. Burillo A, Pedro-Botet ML, Bouza E. 2017. Microbiology and Epidemiology of Legionnaire’s Disease. Infect Dis Clin North Am 31:7–27.

2. Cunha BA, Burillo A, Bouza E. 2016. Legionnaires’ disease. Lancet 387:376–385.

3. Den Boer JW, Nijhof J, Friesema I. 2006. Risk factors for sporadic community-acquired Legionnaires’ disease. A 3-year national case-control study. Public Health 120:566–71.

4. McDade JE, Shepard CC, Fraser DW, Tsai TR, Redus MA, Dowdle WR. 1977. Legionnaires’ disease: isolation of a bacterium and demonstration of its role in other respiratory disease. N Engl J Med 297:1197–203.

5. Boamah DK, Zhou G, Ensminger AW, O’Connor TJ. 2017. From many hosts, one accidental pathogen: the diverse protozoan hosts of *Legionella*. Front Cell Infect Microbiol 7:477.

6. Gimenez G, Bertelli C, Moliner C, Robert C, Raoult D, Fournier PE, Greub G. 2011. Insight into cross-talk between intra-amoebal pathogens. BMC Genomics 12:542.

7. Horwitz MA. 1983. Formation of a novel phagosome by the Legionnaires’ disease bacterium (Legionella pneumophila) in human monocytes. J Exp Med 158:1319–31.

8. Horwitz MA, Maxfield FR. 1984. Legionella pneumophila inhibits acidification of its phagosome in human monocytes. J Cell Biol 99:1936–43.

9. Derre I, Isberg RR. 2004. Legionella pneumophila replication vacuole formation involves rapid recruitment of proteins of the early secretory system. Infect Immun 72:3048–53.

10. Kagan JC, Stein MP, Pypaert M, Roy CR. 2004. Legionella subvert the functions of Rab1 and Sec22b to create a replicative organelle. J Exp Med 199:1201–11.

11. Kagan JC, Roy CR. 2002. Legionella phagosomes intercept vesicular traffic from endoplasmic reticulum exit sites. Nat Cell Biol 4:945–54.

12. Tilney LG, Harb OS, Connelly PS, Robinson CG, Roy CR. 2001. How the parasitic bacterium Legionella pneumophila modifies its phagosome and transforms it into rough ER: implications for conversion of plasma membrane to the ER membrane. J Cell Sci 114:4637–50.

13. Kotewicz KM, Ramabhadran V, Sjoblom N, Vogel JP, Haenssler E, Zhang M, Behringer J, Scheck RA, Isberg RR. 2017. A Single Legionella Effector Catalyzes a Multistep Ubiquitination Pathway to Rearrange Tubular Endoplasmic Reticulum for Replication. Cell Host Microbe 21:169–181.

14. Segal G, Purcell M, Shuman HA. 1998. Host cell killing and bacterial conjugation require overlapping sets of genes within a 22-kb region of the *Legionella pneumophila* genome. Proc Natl Acad Sci USA 95:1669–1674.

15. Vogel JP, Andrews HL, Wong SK, Isberg RR. 1998. Conjugative transfer by the virulence system of Legionella pneumophila. Science 279:873–6.

16. Luo ZQ, Isberg RR. 2004. Multiple substrates of the Legionella pneumophila Dot/Icm system identified by interbacterial protein transfer. Proc Natl Acad Sci U S A 101:841–6.

17. Huang L, Boyd D, Amyot WM, Hempstead AD, Luo ZQ, O’Connor TJ, Chen C, Machner M, Montminy T, Isberg RR. 2010. The E Block motif is associated with *Legionella pneumophila* translocated substrates. Cell Microbiol 13:227–45.

18. Zhu W, Banga S, Tan Y, Zheng C, Stephenson R, Gately J, Luo ZQ. 2011. Comprehensive identification of protein substrates of the Dot/Icm type IV transporter of Legionella pneumophila. PLoS One 6:e17638.

19. Burstein D, Zusman T, Degtyar E, Viner R, Segal G, Pupko T. 2009. Genome-scale identification of Legionella pneumophila effectors using a machine learning approach. PLoS Pathog 5:e1000508.

20. Burstein D, Amaro F, Zusman T, Lifshitz Z, Cohen O, Gilbert AJ, Pupko T, Shuman HA, Segal G. 2016. Genomic analysis of 38 *Legionella* species identifies large and diverse effector repertoires. Nature Genetics 48:167–175.

21. Gomez-Valero L, Rusniok C, Carson D, Mondino S, Perez-Cobas AE, Rolando M, Pasricha S, Reuter S, Demirtas J, Crumbach J, Descorps-Declere S, Hartland EL, Jarraud S, Dougan G, Schroeder GN, Frankel G, Buchrieser C. 2019. More than 18,000 effectors in the *Legionella* genus genome provide multiple, independent combinations for replication in human cells. Proc Natl Acad Sci USA 116:2265–2273.

22. O’Connor TJ, Boyd D, Dorer MS, Isberg RR. 2012. Aggravating genetic interactions allow a solution to redundancy in a bacterial pathogen. Science 338:1440–4.

23. Park JM, Ghosh S, O’Connor TJ. 2020. Combinatorial selection in amoebal hosts drives the evolution of the human pathogen Legionella pneumophila. Nat Microbiol 5:599–609.

24. Neunuebel MR, Chen Y, Gaspar AH, Backlund PS, Jr., Yergey A, Machner MP. 2011. De-AMPylation of the small GTPase Rab1 by the pathogen *Legionella pneumophila*. Science 333:453–456.

25. Kim S, Isberg RR. 2023. The Sde phosphoribosyl-linked ubiquitin transferases protect the Legionella pneumophila vacuole from degradation by the host. Proc Natl Acad Sci U S A 120:e2303942120.

26. Li Z, Dugan AS, Bloomfield G, Skelton J, Ivens A, Losick V, Isberg RR. 2009. The amoebal MAP kinase response to Legionella pneumophila is regulated by DupA. Cell Host Microbe 6:253–67.

27. Shin S, Case CL, Archer KA, Nogueira CV, Kobayashi KS, Flavell RA, Roy CR, Zamboni DS. 2008. Type IV secretion-dependent activation of host MAP kinases induces an increased proinflammatory cytokine response to Legionella pneumophila. PLoS Pathog 4:e1000220.

28. Fontana MF, Banga S, Barry KC, Shen X, Tan Y, Luo ZQ, Vance RE. 2011. Secreted bacterial effectors that inhibit host protein synthesis are critical for induction of the innate immune response to virulent Legionella pneumophila. PLoS Pathog 7:e1001289.

29. Quaile AT, Stogios PJ, Egorova O, Evdokimova E, Valleau D, Nocek B, Kompella PS, Peisajovich S, Yakunin AF, Ensminger AW, Savchenko A. 2018. The Legionella pneumophila effector Ceg4 is a phosphotyrosine phosphatase that attenuates activation of eukaryotic MAPK pathways. J Biol Chem 293:3307–3320.

30. Johnson GL, Lapadat R. 2002. Mitogen-activated protein kinase pathways mediated by ERK, JNK, and p38 protein kinases. Science 298:1911–2.

31. Chang L, Karin M. 2001. Mammalian MAP kinase signalling cascades. Nature 410:37–40.

32. Davis RJ. 2000. Signal transduction by the JNK group of MAP kinases. Cell 103:239–52.

33. Nandi I, Aroeti B. 2023. Mitogen-Activated Protein Kinases (MAPKs) and Enteric Bacterial Pathogens: A Complex Interplay. Int J Mol Sci 24.

34. Losick VP, Haenssler E, Moy MY, Isberg RR. 2010. LnaB: a Legionella pneumophila activator of NF-kappaB. Cell Microbiol 12:1083–97.

35. Iglewicz B, Hoaglin D. 1993. How to Detect and Handle Outliers. *In* Mykytka EF (ed), The ASQC Basic References in Quality Control: Statistical Techniques, vol 16. American Society for Quality Control.

36. Bhogaraju S, Kalayil S, Liu Y, Bonn F, Colby T, Matic I, Dikic I. 2016. Phosphoribosylation of Ubiquitin Promotes Serine Ubiquitination and Impairs Conventional Ubiquitination. Cell 167:1636–1649 e13.

37. Qiu J, Sheedlo MJ, Yu K, Tan Y, Nakayasu ES, Das C, Liu X, Luo ZQ. 2016. Ubiquitination independent of E1 and E2 enzymes by bacterial effectors. Nature 533:120–124.

38. Zhang M, McEwen JM, Sjoblom NM, Kotewicz KM, Isberg RR, Scheck RA. 2021. Members of the Legionella pneumophila Sde family target tyrosine residues for phosphoribosyl-linked ubiquitination. RSC Chem Biol 2:1509–1519.

39. Mukherjee S, Liu X, Arasaki K, McDonough J, Galan JE, Roy CR. 2011. Modulation of Rab GTPase function by a protein phosphocholine transferase. Nature 477:103–6.

40. Gaspar AH, Machner MP. 2014. VipD is a Rab5-activated phospholipase A1 that protects *Legionella pneumophila* from endosomal fusion. Proc Natl Acad Sci U S A 111:4560–5.

41. Ku B, Lee KH, Park WS, Yang CS, Ge J, Lee SG, Cha SS, Shao F, Heo WD, Jung JU, Oh BH. 2012. VipD of *Legionella pneumophila* targets activated Rab5 and Rab22 to interfere with endosomal trafficking in macrophages. PLoS Pathog 8:e1003082.

42. VanRheenen SM, Luo ZQ, O’Connor T, Isberg RR. 2006. Members of a *Legionella pneumophila* family of proteins with ExoU (phospholipase A) active sites are translocated to target cells. Infect Immun 74:3597–606.

43. Heidtman M, Chen EJ, Moy MY, Isberg RR. 2009. Large-scale identification of Legionella pneumophila Dot/Icm substrates that modulate host cell vesicle trafficking pathways. Cell Microbiol 11:230–48.

44. Hohmann S. 2002. Osmotic stress signaling and osmoadaptation in yeasts. Microbiol Mol Biol Rev 66:300–72.

45. Tatebayashi K, Yamamoto K, Tanaka K, Tomida T, Maruoka T, Kasukawa E, Saito H. 2006. Adaptor functions of Cdc42, Ste50, and Sho1 in the yeast osmoregulatory HOG MAPK pathway. EMBO J 25:3033-44.

46. Rep M, Krantz M, Thevelein JM, Hohmann S. 2000. The transcriptional response of Saccharomyces cerevisiae to osmotic shock. Hot1p and Msn2p/Msn4p are required for the induction of subsets of high osmolarity glycerol pathway-dependent genes. J Biol Chem 275:8290–300.

47. O’Rourke SM, Herskowitz I. 1998. The Hog1 MAPK prevents cross talk between the HOG and pheromone response MAPK pathways in Saccharomyces cerevisiae. Genes Dev 12:2874–86.

48. Tamas MJ, Luyten K, Sutherland FC, Hernandez A, Albertyn J, Valadi H, Li H, Prior BA, Kilian SG, Ramos J, Gustafsson L, Thevelein JM, Hohmann S. 1999. Fps1p controls the accumulation and release of the compatible solute glycerol in yeast osmoregulation. Mol Microbiol 31:1087–104.

49. de Felipe KS, Pampou S, Jovanovic OS, Pericone CD, Ye SF, Kalachikov S, Shuman HA. 2005. Evidence for acquisition of Legionella type IV secretion substrates via interdomain horizontal gene transfer. J Bacteriol 187:7716–26.

50. Pan X, Luhrmann A, Satoh A, Laskowski-Arce MA, Roy CR. 2008. Ankyrin repeat proteins comprise a diverse family of bacterial type IV effectors. Science 320:1651–4.

51. Jung US, Sobering AK, Romeo MJ, Levin DE. 2002. Regulation of the yeast Rlm1 transcription factor by the Mpk1 cell wall integrity MAP kinase. Mol Microbiol 46:781–789.

52. Levin DE. 2011. Regulation of cell wall biogenesis in *Saccharomyces cerevisiae*: the cell wall integrity signaling pathway. Genetics 189:1145–1175.

53. de Nadal E, Posas F. 2022. The HOG pathway and the regulation of osmoadaptive responses in yeast. FEMS Yeast Res 22.

54. Dowen RH, Engel JL, Shao F, Ecker JR, Dixon JE. 2009. A family of bacterial cysteine protease type III effectors utilizes acylation-dependent and -independent strategies to localize to plasma membranes. J Biol Chem 284:15867–79.

55. Shao F, Merritt PM, Bao Z, Innes RW, Dixon JE. 2002. A Yersinia effector and a Pseudomonas avirulence protein define a family of cysteine proteases functioning in bacterial pathogenesis. Cell 109:575–88.

56. Holz C, Lueking A, Bovekamp L, Gutjahr C, Bolotina N, Lehrach H, Cahill DJ. 2001. A human cDNA expression library in yeast enriched for open reading frames. Genome Res 11:1730–5.

57. Urbanus ML, Zheng TM, Khusnutdinova AN, Banh D, Mount HOC, Gupta A, Stogios PJ, Savchenko A, Isberg RR, Yakunin AF, Ensminger AW. 2024. A random mutagenesis screen enriched for missense mutations in bacterial effector proteins. bioRxiv doi:10.1101/2024.03.14.585084:2024.03.14.585084.

58. Jumper J, Evans R, Pritzel A, Green T, Figurnov M, Ronneberger O, Tunyasuvunakool K, Bates R, Zidek A, Potapenko A, Bridgland A, Meyer C, Kohl SAA, Ballard AJ, Cowie A, Romera-Paredes B, Nikolov S, Jain R, Adler J, Back T, Petersen S, Reiman D, Clancy E, Zielinski M, Steinegger M, Pacholska M, Berghammer T, Bodenstein S, Silver D, Vinyals O, Senior AW, Kavukcuoglu K, Kohli P, Hassabis D. 2021. Highly accurate protein structure prediction with AlphaFold. Nature 596:583–589.

59. Mirdita M, Schutze K, Moriwaki Y, Heo L, Ovchinnikov S, Steinegger M. 2022. ColabFold: making protein folding accessible to all. Nat Methods 19:679–682.

60. Lucas M, Gaspar AH, Pallara C, Rojas AL, Fernandez-Recio J, Machner MP, Hierro A. 2014. Structural basis for the recruitment and activation of the *Legionella* phospholipase VipD by the host GTPase Rab5. Proc Natl Acad Sci U S A 111:E3514–23.

61. Ge J, Xu H, Li T, Zhou Y, Zhang Z, Li S, Liu L, Shao F. 2009. A Legionella type IV effector activates the NF-kappaB pathway by phosphorylating the IkappaB family of inhibitors. Proc Natl Acad Sci U S A 106:13725–30.

62. Fontana MF, Shin S, Vance RE. 2012. Activation of host mitogen-activated protein kinases by secreted Legionella pneumophila effectors that inhibit host protein translation. Infect Immun 80:3570–5.

63. Cargnello M, Roux PP. 2011. Activation and function of the MAPKs and their substrates, the MAPK-activated protein kinases. Microbiol Mol Biol Rev 75:50–83.

64. de Felipe KS, Glover RT, Charpentier X, Anderson OR, Reyes M, Pericone CD, Shuman HA. 2008. *Legionella* eukaryotic-like type IV substrates interfere with organelle trafficking. PLoS Pathog 4:e1000117.

65. Mukherjee S, Keitany G, Li Y, Wang Y, Ball HL, Goldsmith EJ, Orth K. 2006. Yersinia YopJ acetylates and inhibits kinase activation by blocking phosphorylation. Science 312:1211–4.

66. Zhu M, Shao F, Innes RW, Dixon JE, Xu Z. 2004. The crystal structure of Pseudomonas avirulence protein AvrPphB: a papain-like fold with a distinct substrate-binding site. Proc Natl Acad Sci U S A 101:302–7.

67. Shao F, Golstein C, Ade J, Stoutemyer M, Dixon JE, Innes RW. 2003. Cleavage of Arabidopsis PBS1 by a bacterial type III effector. Science 301:1230–3.

68. Nachmias N, Zusman T, Segal G. 2019. Study of *Legionella* effector domains revealed novel and prevalent phosphatidylinositol 3-phosphate binding domains. Infect Immun 87:e00153–19.

69. Jank T, Bohmer KE, Tzivelekidis T, Schwan C, Belyi Y, Aktories K. 2012. Domain organization of *Legionella* effector SetA. Cell Microbiol 14:852–68.

70. Pike CM, Boyer-Andersen R, Kinch LN, Caplan JL, Neunuebel MR. 2019. *Legionella* effector RavD binds phosphatidylinositol-3-phosphate and helps suppress endolysosomal maturation of the *Legionella*-containing vacuole. J Biol Chem 294:6405–6415.

71. Weber SS, Ragaz C, Hilbi H. 2009. The inositol polyphosphate 5-phosphatase OCRL1 restricts intracellular growth of *Legionella*, localizes to the replicative vacuole and binds to the bacterial effector LpnE. Cell Microbiol 11:442–460.

72. Ozhelvaci F, Steczkiewicz K. 2023. Identification and classification of papain-like cysteine proteinases. J Biol Chem 299:104801.

73. Mukherjee S, Keitany G, Li Y, Wang Y, Ball HL, Goldsmith EJ, Orth K. 2006. YopJ acetylates and inhibits kinase activation by blocking phosphorylation. Science 312:1211–4.

74. Joseph AM, Shames SR. 2021. Affecting the Effectors: Regulation of Legionella pneumophila Effector Function by Metaeffectors. Pathogens 10.

75. Tan Y, Arnold RJ, Luo ZQ. 2011. Legionella pneumophila regulates the small GTPase Rab1 activity by reversible phosphorylcholination. Proc Natl Acad Sci U S A 108:21212–7.

76. Campanacci V, Mukherjee S, Roy CR, Cherfils J. 2013. Structure of the Legionella effector AnkX reveals the mechanism of phosphocholine transfer by the FIC domain. EMBO J 32:1469–77.

77. Chung N, Zhang XD, Kreamer A, Locco L, Kuan PF, Bartz S, Linsley PS, Ferrer M, Strulovici B. 2008. Median absolute deviation to improve hit selection for genome-scale RNAi screens. J Biomol Screen 13:149–58.

78. Gietz RD, Woods RA. 2002. Transformation of yeast by lithium acetate/single-stranded carrier DNA/polyethylene glycol method. Methods Enzymol 350:87–96.

79. Jansen G, Wu C, Schade B, Thomas DY, Whiteway M. 2005. Drag&Drop cloning in yeast. Gene 344:43–51.

80. Ho SN, Hunt HD, Horton RM, Pullen JK, Pease LR. 1989. Site-directed mutagenesis by overlap extension using the polymerase chain reaction. Gene 77:51–59.

81. Amyot WM, deJesus D, Isberg RR. 2013. Poison domains block transit of translocated substrates via the Legionella pneumophila Icm/Dot system. Infect Immun 81:3239–52.

82. Wang J, Youkharibache P, Zhang D, Lanczycki CJ, Geer RC, Madej T, Phan L, Ward M, Lu S, Marchler GH, Wang Y, Bryant SH, Geer LY, Marchler-Bauer A. 2020. iCn3D, a web-based 3D viewer for sharing 1D/2D/3D representations of biomolecular structures. Bioinformatics 36:131–135.

