## Supplementary Fig. S1 for "Genetic evidence for a regulated cysteine protease catalytic triad in LegA7, a *Legionella pneumophila* protein that impinges on a stress response pathway"

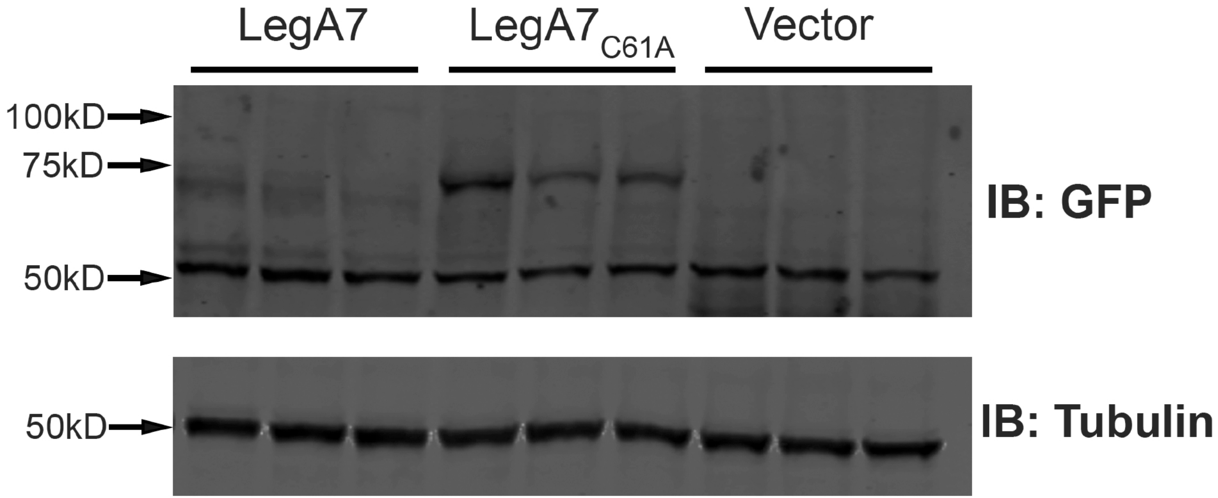


**Supplemental Fig. S1. Mutation in putative catalytic residue C61 reduces degradation of LegA7.** HEK293T were transfected with the noted plasmids and 40 hrs after transfection, the cultures were extracted, gel fractionated and probed with noted antibodies (Materials and Methods). Shown are three independent transfections with each plasmid. LegA7: GFP-LegA7 derivative described in text (Materials and Methods); $LegA7_{C61A}$: GFP-LegA7 having Cys->Ala mutation in predicted catalytic residue C61.
