## Supplementary Fig. S2 for "Genetic evidence for a regulated cysteine protease catalytic triad in LegA7, a *Legionella pneumophila* protein that impinges on a stress response pathway"

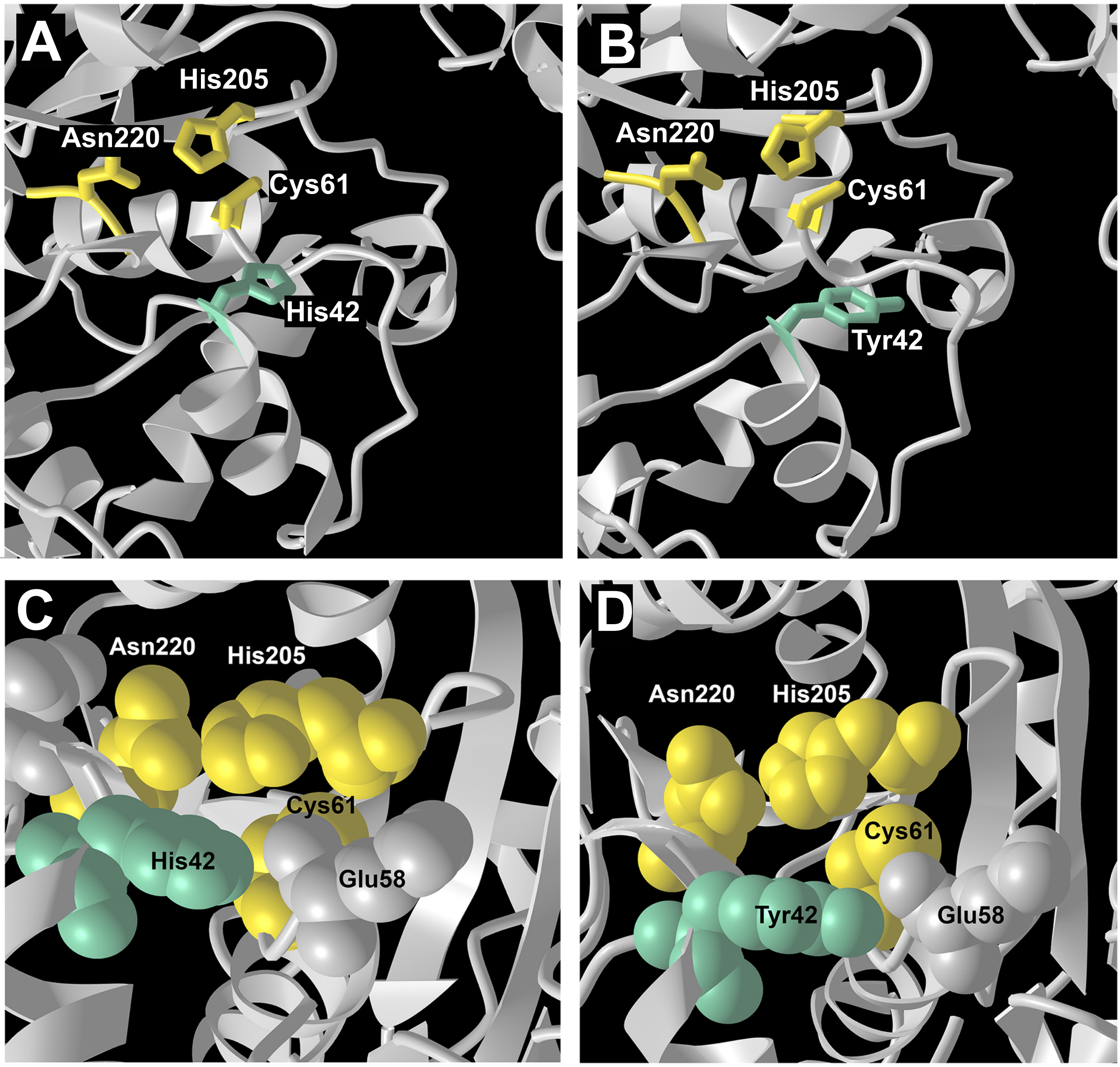


**Fig. S2. Modeling of H42Y mutation places the Tyr sidechain proximal to the LegA7 cysteine protease catalytic site.** Shown is the LegA7 catalytic triad (Cys61, His205, Asn220) displayed as yellow sidechains, with the nearby His42 (green) or the Tyr42 replacement. (A, B) Stick figures of relevant sidechains for WT (A) and Tyr42 mutation (B). (C,D) Space filling models of relevant sidechains for WT (C) and Tyr42 mutation (D).
