## Supplementary FIg. S3 for "Genetic evidence for a regulated cysteine protease catalytic triad in LegA7, a *Legionella pneumophila* protein that impinges on a stress response pathway"

\> Δank-1(331-454)_deletion endpoints (sense)
CGACAACGACTTAGTGATGATGAGGAGAGATTGGAAAAAGAA
R Q R L S D D E E R L E K E

\>Δank-1(331-454)_translation
**MIYVLICIMLVCGLVVYQLMPKLLSYSTSLISKKQDTSELTHDGITNVLTQLGHPKFEGVCYGFTLNWALAVAQGKESFFYRQLH**
**HLRTHQFGLPETLQQIKEKKERNQSLSKDEKIIETLPQLGKKICIAQDPLQYKEKYKKLVWQPDINSILKAINADSSVAKHIFYK**
**THSFLNQDEATEYLELLKRTGIREDVAVIISTADHAMGFKLAGNVWRFININDLYQQDKNKPYFEFSSRNLVKELYRVCAENLQG**
**SRLTVNTDFVSVNPEEKLSRALQNLFPVFPVRTKTSYPERLAFFSMAATQGDMNSVKKCIHSGWSIFSRQRLSDD**SPILTAIYLG
RRDVVRAMLSTSRHRVNQKRKSDSSTLLHIACRYGGSGIVEDLLNIRGIKIDPRDSKGRTPLMYACKKSVVTEDRKLFNLLFAKG
ASLSIKDNDGLTALDHALKNEHTLAIQMI**EERLEKEACAQENSTSRRFKFSETKGTLFQRGVKTISYQSQQPRFGMK**

\> Δank-2(290-454)_ deletion endpoints(sense)
TTTCCAGTCAGAACCAAAGAGGAGAGATTGGAAAAAGAA
F P V R T K E E R L E K E

\> Δank-2(290-454)_translation
**MIYVLICIMLVCGLVVYQLMPKLLSYSTSLISKKQDTSELTHDGITNVLTQLGHPKFEGVCYGFTLNWALAVAQGKESFFYRQLH**
**HLRTHQFGLPETLQQIKEKKERNQSLSKDEKIIETLPQLGKKICIAQDPLQYKEKYKKLVWQPDINSILKAINADSSVAKHIFYK**
**THSFLNQDEATEYLELLKRTGIREDVAVIISTADHAMGFKLAGNVWRFININDLYQQDKNKPYFEFSSRNLVKELYRVCAENLQG**
**SRLTVNTDFVSVNPEEKLSRALQNLFPVFPVRTK**TSYPERLAFFSMAATQGDMNSVKKCIHSGWSIFSRQRLSDDSPILTAIYLG
RRDVVRAMLSTSRHRVNQKRKSDSSTLLHIACRYGGSGIVEDLLNIRGIKIDPRDSKGRTPLMYACKKSVVTEDRKLFNLLFAKG
ASLSIKDNDGLTALDHALKNEHTLAIQMI**EERLEKEACAQENSTSRRFKFSETKGTLFQRGVKTISYQSQQPRFGMK**

**\>** ΔN-ank-1(264-361)_deletion endpoints(sense)
CGATTGACCGTGAATACCGATAGCGACTCTTCTACTTTGCTC
R L T V N T D S D S S T L L

\> ΔN-ank-1(264-361)_translation
**MIYVLICIMLVCGLVVYQLMPKLLSYSTSLISKKQDTSELTHDGITNVLTQLGHPKFEGVCYGFTLNWALAVAQGKESFFYRQLH**
**HLRTHQFGLPETLQQIKEKKERNQSLSKDEKIIETLPQLGKKICIAQDPLQYKEKYKKLVWQPDINSILKAINADSSVAKHIFYK**
**THSFLNQDEATEYLELLKRTGIREDVAVIISTADHAMGFKLAGNVWRFININDLYQQDKNKPYFEFSSRNLVKELYRVCAENLQG**
**SRLTVNTD**FVSVNPEEKLSRALQNLFPVFPVRTKTSYPERLAFFSMAATQGDMNSVKKCIHSGWSIFSRQRLSDDSPILTAIYLG
RRDVVRAMLSTSRHRVNQKRK**SDSSTLLHIACRYGGSGIVEDLLNIRGIKIDPRDSKGRTPLMYACKKSVVTEDRKLFNLLFAKG**
**ASLSIKDNDGLTALDHALKNEHTLAIQMIEERLEKEACAQENSTSRRFKFSETKGTLFQRGVKTISYQSQQPRFGMK**

**\>** ΔN-ank-2(290-361)_deletion endpoints(sense)
GTGTTTCCAGTCAGAACCAAAAGCGACTCTTCTACTTTGCTC
V F P V R T K S D S S T L L

\> ΔN-ank-2(290-361)_translation
**MIYVLICIMLVCGLVVYQLMPKLLSYSTSLISKKQDTSELTHDGITNVLTQLGHPKFEGVCYGFTLNWALAVAQGKESFFYRQLH**
**HLRTHQFGLPETLQQIKEKKERNQSLSKDEKIIETLPQLGKKICIAQDPLQYKEKYKKLVWQPDINSILKAINADSSVAKHIFYK**
**THSFLNQDEATEYLELLKRTGIREDVAVIISTADHAMGFKLAGNVWRFININDLYQQDKNKPYFEFSSRNLVKELYRVCAENLQG**
**SRLTVNTDFVSVNPEEKLSRALQNLFPVFPVRTK**TSYPERLAFFSMAATQGDMNSVKKCIHSGWSIFSRQRLSDDSPILTAIYLG
RRDVVRAMLSTSRHRVNQKRK**SDSSTLLHIACRYGGSGIVEDLLNIRGIKIDPRDSKGRTPLMYACKKSVVTEDRKLFNLLFAKG**
**ASLSIKDNDGLTALDHALKNEHTLAIQMIEERLEKEACAQENSTSRRFKFSETKGTLFQRGVKTISYQSQQPRFGMK**

**Supplemental Figure S3.** Endpoints of deletions described in Fig. 6. Shown are the DNA sequence endpoints of each deletion mutant described in Fig. 6 and the predicted amino acid sequence of each derivative, based on whole plasmid sequencing (Materials and Methods). Red: nucleotide sequence of 3' end of deletion. Black Bold: amino acid sequences remaining in derivative. Gray: amino acid sequences missing.
